# Tyrosine phosphorylation tunes chemical and thermal sensitivity of TRPV2 ion channel

**DOI:** 10.1101/2022.03.03.482857

**Authors:** Xiaoyi Mo, Peiyuan Pang, Yulin Wang, Dexiang Jiang, Mengyu Zhang, Yang Li, Peiyu Wang, Qizhi Geng, Chang Xie, Hai-Ning Du, Bo Zhong, Dongdong Li, Jing Yao

## Abstract

Transient receptor potential vanilloid 2 (TRPV2) is a multimodal ion channel widely regulating central and peripheral functions. Its important involvement in immune responses has been suggested such as in the macrophages’ phagocytosis process. However, the endogenous signaling cascades controlling the gating of TRPV2 remain to be understood. Here, we report that enhancing tyrosine phosphorylation remarkably alters the chemical and thermal sensitivities of TRPV2 endogenously expressed in rat bone marrow-derived macrophages. We identify that the protein tyrosine kinase JAK1 mediates TRPV2 phosphorylation at the molecular sites Tyr(335), Tyr(471), and Tyr(525). JAK1 phosphorylation is required for maintaining TRPV2 activity and the phagocytic ability of macrophages. We further show that TRPV2 phosphorylation is dynamically balanced by protein tyrosine phosphatase non-receptor type 1 (PTPN1). PTPN1 inhibition increases TRPV2 phosphorylation, further reducing the activation temperature threshold to ∼40 °C. Our data thus unveil an intrinsic mechanism where the phosphorylation/dephosphorylation dynamic balance sets the basal chemical and thermal sensitivity of TRPV2. Targeting this pathway will aid therapeutic interventions in physiopathological contexts.

## Introduction

Transient receptor potential vanilloid 2 (TRPV2) channel is broadly expressed in the body, such as the nervous system (Caterina et al., 1999; Nedungadi et al., 2012), the immune system (Link et al., 2010; Nagasawa et al., 2007), and the muscular system (Peng et al., 2010; Zanou et al., 2015). As a Ca^2+^ permeable polymodal receptor, TRPV2 responds to noxious temperature (> 52 °C) (Caterina et al., 1999), mechanical force (McGahon et al., 2016; Sugio et al., 2017), osmotic swelling (Muraki et al., 2003), and chemical modulators including 2-Aminoethyl diphenylborinate (2-APB) (Hu et al., 2004), cannabinoids (De Petrocellis et al., 2011), probenecid (Bang et al., 2007), tranilast (Iwata et al., 2020) and SKF96365 (Juvin et al., 2007). TRPV2 has been implicated in diverse biological functions including thermal sensation (Caterina et al., 1999), neuronal development (Shibasaki et al., 2010), osmotic- or mechanosensation (Muraki et al., 2003; Sugio et al., 2017), cardiac-structure maintenance (Katanosaka et al., 2014), insulin secretion (Aoyagi et al., 2010), proinflammatory process (Entin-Meer et al., 2017; Yamashiro et al., 2010) and oncogenesis (Siveen et al., 2020). The role of TRPV2 in immune responses has also been suggested (Link et al., 2010; Santoni et al., 2013), such as its regulation of macrophage particle binding and phagocytosis (Link et al., 2010). In mast cells, TRPV2-mediated calcium flux stimulates protein kinase A (PKA)-dependent proinfammation degranulation (Stokes et al., 2004). In addition, early studies have shown that peripheral inflammation and phosphoinositide 3-kinase (PI3K) signaling pathways enhance TRPV2 function by recruiting it onto the plasma membrane (Aoyagi et al., 2010; Shimosato et al., 2005).

At the channel level, our recent study found that the lipid-raft-associated protein flotillin-1 interacts with and sustains the surface expression of the TRPV2 channel (Hu et al., 2021). The use dependence of the TRPV2 channel in heat sensitivity but not agonist sensitivity has also been reported (Liu and Qin, 2016). Recently, the oxidation of TRPV2 on methionine residues was found to activate and sensitize the channel (Fricke et al., 2019). Moreover, the structure of TRPV2 at near-atomic resolution has been determined by cryo-electron microscopy (Huynh et al., 2016; Zubcevic et al., 2016). Despite the functional and structural insights, the endogenous signaling elements that gate TRPV2 activities remain poorly understood.

Here we show that the phosphokinases regulator magnesium (Mg^2+^), exerts an enhancing effect on both the chemical and thermal sensitivity of TRPV2 endogenously expressed in rat bone marrow-derived macrophages. We then provide evidence that Mg^2+^ activates the phosphokinase JAK1 to increase the phosphorylation levels of TRPV2. In contrast, JAK1 inhibition downregulates TRPV2 channel activity, which in accordance reduces the phagocytic ability of macrophages. We have also determined three JAK1 phosphorylation sites, Y335, Y471, and Y525, in TRPV2. Further, we identify that PTPN1 is the tyrosine phosphatase that mediates TRPV2 dephosphorylation. Our data unmask an endogenous signaling cascade where tyrosine phosphorylation homeostasis contributes to setting the sensitivity of TRPV2 to thermal and chemical stimuli. These observations should help to conceive potential therapeutic targeting of TRPV2 in physiological and pathological situations.

## Results

### Mg^2+^ enhances both the chemical and thermal sensitivity of TRPV2

Enriched in cell cytoplasm, Mg^2+^ regulates the function of a variety of ion channels (Antonov and Johnson, 1999; Cao et al., 2014; Lee et al., 2005; Luo et al., 2012; Obukhov and Nowycky, 2005). We, therefore, sought to examine whether TRPV2 activity is sensitive to Mg^2+^. Considering that TRPV2 is abundantly and functionally expressed in macrophages where other types of TRPV channels are barely detectable (Link et al., 2010; Nagasawa et al., 2007), We hence used rat bone marrow-derived macrophages (rBMDMs) as an endogenous cell system to record TRPV2 currents. We found that TRPV2 currents at -60 mV evoked by 0.3 mM 2-APB were slowly but dramatically enhanced in the presence of 5 mM Mg^2+^ (Figure 1A). The pipette solution contained 1 mM adenosine disodium triphosphate (Na_2_ATP). In general, Mg^2+^- potentiated responses typically developed over a period of about 100 s to reach a plateau. The presence of 5 mM Mg^2+^ augmented the peak current amplitudes by ∼19-fold (Figure 1B). Notably, the effect of Mg^2+^ could not be completely washed out and the following response to 0.3 mM 2-APB was somewhat variable but still remained ∼14-fold increase to that before Mg^2+^ treatment (Figure 1A-B). We further recorded the effect of Mg^2+^ on TRPV2 current responses in neurons. TRPV2 channels are predominantly expressed in medium- to large-sized dorsal root ganglia (DRG) neurons that typically express fewer TRPV1 channels (Caterina et al., 1999). As illustrated in Figure 1C-D, we witnessed similar potentiating effects of Mg^2+^ on 2-APB-evoked currents in a small population of DRG neurons, while the lack of TRPV1 expression was confirmed by the absence of responses to capsaicin, indicating these 2-APB-evoked currents were mediated by TRPV2 channels. To further investigate whether the regulatory effect of Mg^2+^ on TRPV2 reflects a channel-inherent mechanism, we performed recordings in a variety of heterologous expression systems including HEK 293T (Figure 1E-F), CHO, Hela, and ND7/23 cells (Figure 1 - figure supplement 1) where TRPV2 was transiently expressed. Indeed, the profound enhancement of TRPV2 activity by Mg^2+^ was observed in all expression cell lines. Next, we asked whether other divalent cations exert similar regulatory effects on TRPV2 currents as Mg^2+^ does. We thus repeated the experiments in TRPV2-expressing HEK 293T cells with different cations including Mn^2+^, Ca^2+^, Ba^2+^, Zn^2+^, Cu^2+^, Ni^2+^, Cd^2+^, and Co^2+^. As shown in Figure 1 - figure supplement 2, among all the tested divalent cations, only Mg^2+^ exhibited a more profound effect on enhancing the TRPV2 channel activity.

**Figure 1.**
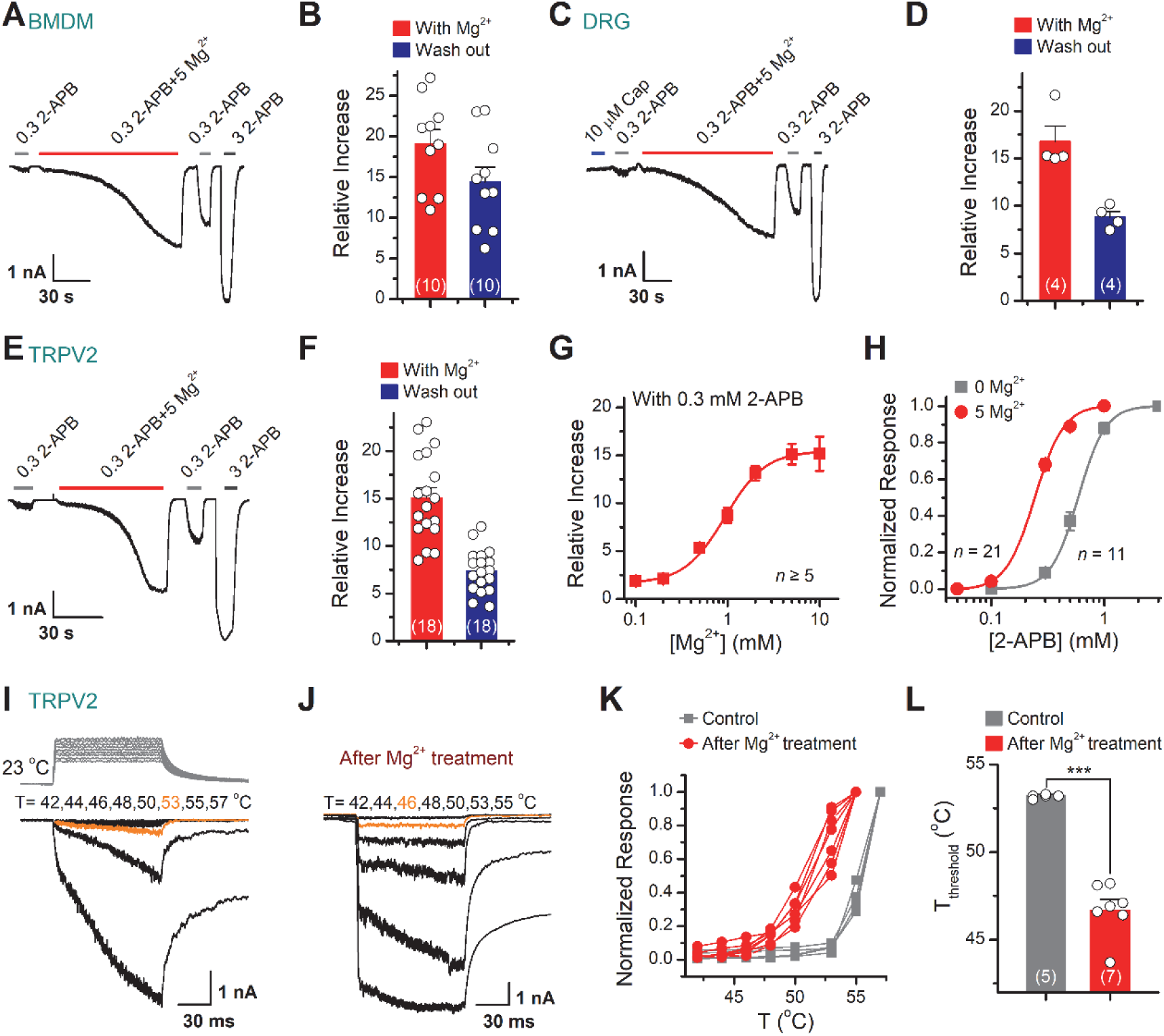
TRPV2 activities are enhanced in the presence of Mg^2+^. (A) Mg^2+^ potentiates 2-APB responses in a representative rat BMDM cell. The cell was exposed to 0.3 mM 2-APB without or with 5 mM Mg^2+^, and 3 mM 2-APB as indicated by the bars. Membrane currents were recorded in whole-cell configuration, and the holding potential was -60 mV. Bars represent duration of drug application. (B) Summary of relative currents evoked by 0.3 mM 2-APB in the presence of 0 or 5 mM Mg^2+^. Numbers of cells are indicated in parentheses. (C) Whole-cell currents at -60 mV in a rat DRG neuron treated with 10 μM Cap, 0.3 mM 2-APB, 0.3 mM 2-APB plus 5 mM Mg^2+^, and 3 mM 2-APB. (D) Summary of relative currents elicited by without or with 5 mM Mg^2+^. (E-F) Parallel whole-cell recordings in TRPV2-expressing HEK 293T cells and the relative changes caused by Mg^2+^. (G) Dose dependence of Mg^2+^ effects on 2-APB response (0.3 mM). The solid line represents a fit by Hill’s equation with EC_50_ = 0.94 ± 0.04 mM and n_H_ = 2.0 ± 0.2 (*n* ≥ 5). (H) Dose-response curves of 2-APB for activation of TRPV2 in the presence of 0 or 5 mM Mg^2+^. The solid lines corresponds to Hill’s equation with EC_50_ = 0.59 ± 0.01 mM and n_H_ = 3.6 ± 0.1 for 0 Mg^2+^ (*n* = 11); and EC_50_ = 0.24 ± 0.01 mM and n_H_ = 3.4 ± 0.1 for application of 5 mM Mg^2+^ (*n* = 21). (I-J) Effects of Mg^2+^ on temperature dependence. Representative responses to a family of temperature pulses for TRPV2-expressing HEK293T cells under control conditions or pretreated with 5 mM Mg^2+^. Temperature pulses stepped from room temperature generated by laser irradiation were 100 ms long and had a rise time of 2 ms. The threshold temperature for heat activation of TRPV2 was determined as the temperature at which ∼10% of its maximum response was induced. (K) Temperature-dependent response curves were measured from the maximal currents at the end of temperature steps. Each curve indicates measurements from an individual cell. (L) Comparison of temperature thresholds for activation of TRPV2. Different symbols represent individual data points. The mean temperature thresholds (T_threshold_) were 53.2 ± 0.1 °C (*n* = 5) for control, and 46.7 ± 0.6 °C (*n* = 7) for post-treatment with 5 mM Mg^2+^. *P* = 2.58E-6 by unpaired student *t*-test. Error bars indicate SEM.

To further characterize the regulatory effects of Mg^2+^ on TRPV2 activity, whole-cell currents were elicited by local perfusion of 0.3 mM 2-APB with varied concentrations of Mg^2+^ ranging from 0.1 to 10 mM. Mg^2+^ was effective above 0.1 mM and remained effective up to 10 mM with a half-maximal concentration of 0.94 ± 0.04 mM (Figure 1G). In addition, the EC_50_ of 2-APB on TRPV2 activation was shifted to 0.24 ± 0.01 mM from 0.59 ± 0.01 mM in the presence of 5 mM Mg^2+^ (Figure 1H).

TRPV2 is a member of the temperature-sensitive ion channel. Therefore, we examined the effect of Mg^2+^ on TRPV2 thermosensitivity using laser irradiation-based temperature controlling and whole-cell recording (Yao et al., 2009). HEK 293T cells expressing TRPV2 were held at −60 mV when the temperature jumps were delivered (Figure 1I, inset). The above experiments showed that the enhanced effect of Mg^2+^ on TRPV2 channel requires long-term continuous treatment, however, prolonged high temperature stimulation incurs excessive thermal stress and leads to the instability of whole-cell recordings. For such a reason, we first sensitized the TRPV2 channel by stimulating the cells with the combination of 0.3 mM 2-APB and 5 mM Mg^2+^, and then immediately applied the temperature pulses to the same cell right after completely washout 2-APB by bath solution. As illustrated in Figure 1I-J, the pretreatment with Mg^2+^ evidently lowered the temperature threshold in TRPV2 activation. Plotting the relative responses revealed that Mg^2+^ caused an apparent left-shift of the temperature dependence curve of TRPV2 (Figure 1K), with the activation temperature threshold being lowered by ∼6 °C (Figure 1L). Together, these results indicate that Mg^2+^ enhances both the chemical and thermal responses of the TRPV2 ion channel.

### Mg^2+^ potentiates TRPV2 activation via an indirect intracellular pathway

To identify whether Mg^2+^ directly activates TRPV2, we recorded its currents in HEK 293T cells using whole-cell patch-clamp in the presence of various concentrations of Mg^2+^ (Figure 2A). We observed that even 100 mM Mg^2+^ did not induce any detectable current (Figure 2A-B), indicating that extracellular Mg^2+^ cannot directly activate TRPV2 channels. Likely, Mg^2+^ enhances TRPV2 activation via an intracellular mechanism. Thus, extracellularly applied Mg^2+^ might need to permeate into cell cytosol through the activated channel. Typically, glutamate residues (E) and aspartate residues (D) of TRPV channels control the permeation of divalent cations. For instance, the TRPV1-D646N/E648Q/E651Q mutant impairs the Ca^2+^ permeability (Samways and Egan, 2011). To probe the mechanism of Mg^2+^-mediated enhancement of TRPV2 activity, we first mutated the equivalent residues, E609/E614 in TRPV2 to glutamine (Q) to impair its divalent cation permeability, which was verified by Ca^2+^ imaging showing a decreased Ca^2+^ influx (Figure 2 - figure supplement 1), and then examined whether the double mutation could alter the Mg^2+^ effect (Figure 2C). As shown in Figure 2D, reducing Mg^2+^ entry indeed eliminated its enhancing effect on TRPV2 whole-cell currents evoked by 2-APB. As corroboration, chelating intracellular Mg^2+^ with 20 mM EDTA delivered through patch pipette also abolished the enhancement effect (Figure 2E-F).

**Figure 2.**
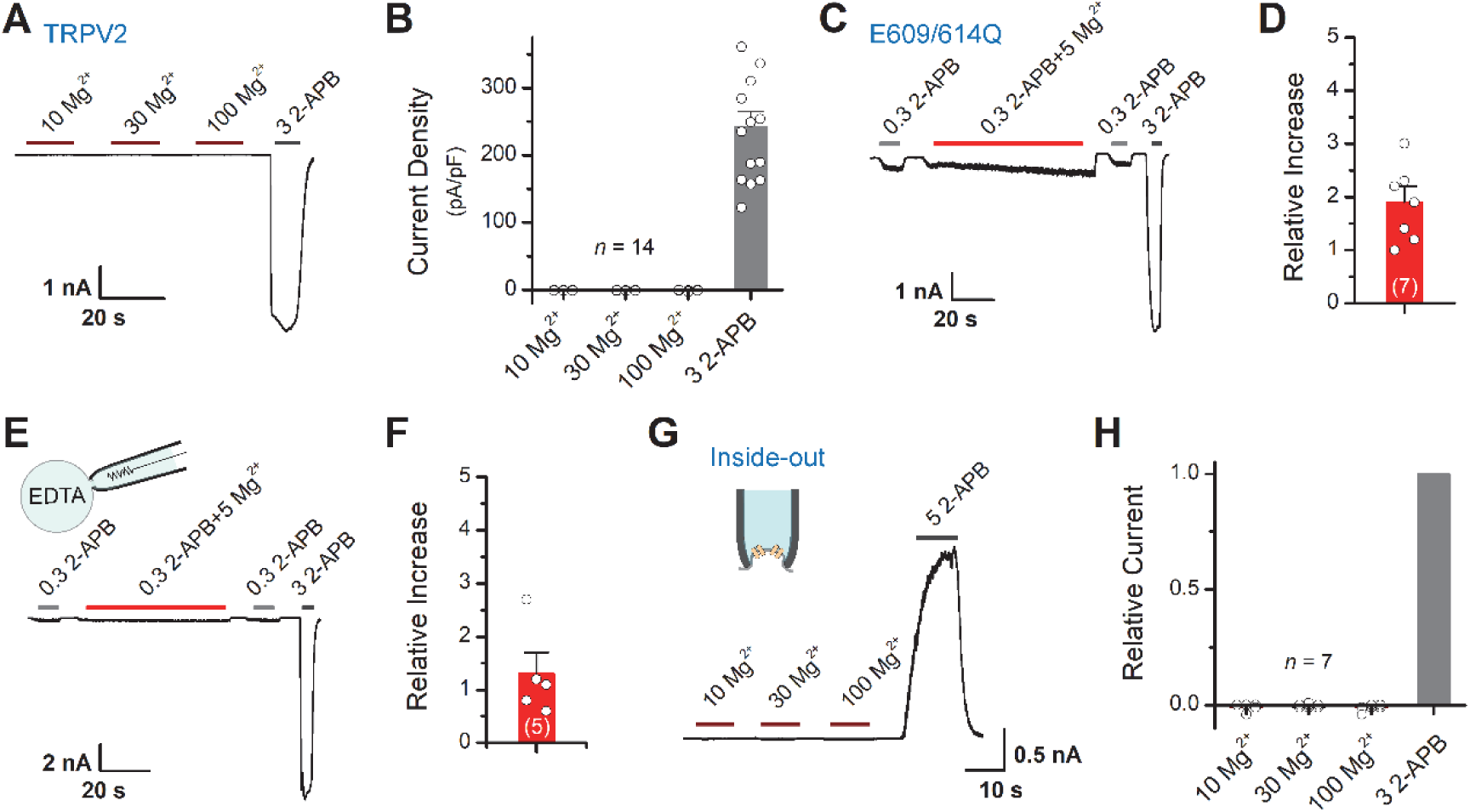
Mg^2+^ has an indirect effect on TRPV2 channels. (A) High concentrations of Mg^2+^ have no direct effect on TRPV2 channels from the extracellular side. Representative whole-cell currents at -60 mV in a TRPV2-expressing HEK 293T cells consecutively treated with 10, 30, 100 mM Mg^2+^ and 3 mM 2-APB. (B) Comparison of current density evoked by different concentrations of Mg^2+^ and 3 mM 2-APB. (C) Representative whole-cell recordings showing that Mg^2+^ failed to potentiate TRPV2(E609Q/E614Q) even though the response to 2-APB was retained. (D) Summary of relative currents elicited by the combination of 0.3 mM 2-APB and 5 mM Mg^2+^ versus 0.3 mM 2-APB. (E) Whole-cell recordings from TRPV2-expressing HEK293T cells showing the response to 0.3 mM 2-APB, 0.3 mM 2-APB plus 5 mM Mg^2+^, and 3 mM 2-APB. Note the pipette solution contained 20 mM EDTA. (F) Average plot of the relative changes. (G) Current traces recorded in inside-out configuration evoked by different concentrations of Mg^2+^ and 5 mM 2-APB. (H) Summary plot of relative currents elicited by 10, 30, 100 mM Mg^2+^ and 3 mM 2-APB. Error bars indicate SEM.

The above results suggest that the enhancing effect of Mg^2+^ on TRPV2 activation takes place on the intracellular side. We then performed inside-out patch-clamp to examine whether Mg^2+^ directly activates TRPV2 from the intracellular side (Figure 2G). Akin to extracellular application, even 100 mM Mg^2+^ did not induce any detectable current from the intracellular side (Figure 2H). Together, our results suggest that the potentiation effect of Mg^2+^ on TRPV2 activation relies on an indirect intracellular mechanism.

### JAK1-mediated tyrosine phosphorylation regulates TRPV2 sensitivity

Previous studies suggest that some stimuli, like insulin, recruit TRPV2 to the plasma membrane to increase the whole-cell response (Hisanaga et al., 2009; Kanzaki et al., 1999; Nagasawa et al., 2007). To verify whether Mg^2+^ solicits similar mechanisms, we compared the saturation currents evoked by a high dose of 2-APB (3 mM) before and after Mg^2+^ treatment. Our data displayed that subsequent to Mg^2+^ application, though the currents evoked by sub-saturation doses of 2-APB were well potentiated, there was no significant change in the maximum saturation currents (Figure 3A). This observation indicates that Mg^2+^ does not alter the expression level of TRPV2 at the plasma membrane.

**Figure 3.**
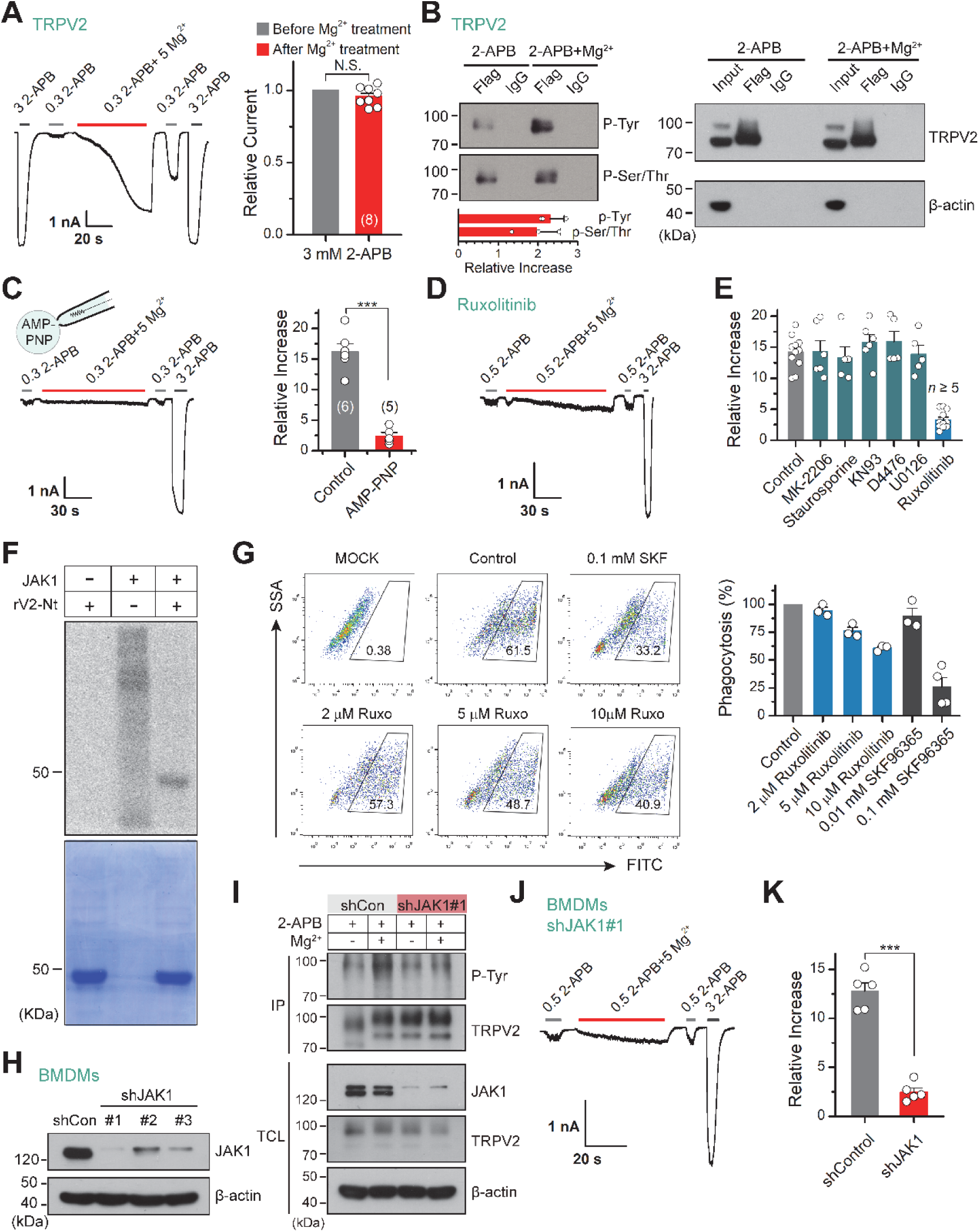
Tyrosine phosphokinase JAK1 upregulates channel activity via phosphorylation of TRPV2. (A) Representative whole-cell recordings from TRPV2-expressing HEK 293T cells showing the responses to 3 mM 2-APB before and after the treatment by 0.3 mM 2-APB plus 5 mM Mg^2+^ (*Left*). Average peak responses to 3 mM 2-APB before and after Mg^2+^ application (*Right*). The holding potential was -60 mV. *P* = 0.12 by one-sample *t*-test. (B) Tyrosine phosphorylation and serine/threonine phosphorylation of immunoprecipitated TRPV2-Flag transiently transfected in HEK293T cells in the absence and presence of 5 mM Mg^2+^ were determined by immunoblotting with anti-phosphotyrosine antibody (pTyr) and anti-Phospho-(Ser/Thr) Phe antibody (pSer/Thr). *Inset*, Protein amounts of tyrosine-phosphorylated or serine/threonine-phosphorylated immunoprecipitated TRPV2 proteins were quantified, and phospho-Tyr TRPV2/total TRPV2 and phospho-Ser/Thr TRPV2/total TRPV2 were calculated from at least three independent experiments. Error bars indicate SD. (C) *Left*, representative whole-cell currents at -60 mV in a TRPV2 –expressing HEK 293T cell treated with 0.3 mM 2- APB, 0.3 mM 2-APB plus 5 mM Mg^2+^ and 3 mM 2-APB. The pipette solution contained ATP nonhydrolyzable analog adenylyl imidodiphosphate (AMP-PNP). *Right*, summary of relative changes under different conditions. *P* = 9.29E-6 by unpaired student *t*-test. (D) Whole-cell currents in response to 2-APB under inhibition of JAK1 by Ruxolitinib. (E) Summary plot of Mg^2+^ effects on TRPV2 currents under the various conditions. (F) *In vitro* kinase assay with [^32^P]-γ-ATP, tyrosine kinase JAK1, and recombinant his-tagged rat TRPV2 N-terminus. Phosphorylation signals were detected by autoradiography. Loading amount of different TRPV2 proteins was accessed by coomassie blue staining. (G) Flow cytometry analysis for phagocytosis. Flow cytometry analysis was employed to determine the phagocytosed level of green fluorescent protein (GFP)-expressing Escherichia coli (GFP E. coli) by BMDMs treated with varying concentrations of Ruxolitinib or SKF96365. Bar graph displaying the effects on phagocytosis under different conditions. (H) Immunoblot analysis (with anti-JAK1 or anti-β actin) of BMDM cells transfected for 72 h with JAK-1-targeting shRNA (shJAK1#1, shJAK1#2 and shJAK1#3) or shCon to test knockdown efficiency of shRNA. (I) Western blot analysis of the tyrosine phosphorylation levels of TRPV2 in BMDM cells transfected with shJAK1#3 or shCon for 72 h in the absence and presence of Mg^2+^, respectively. (J) Whole-cell recordings in BMDM cells transfected with shJAK1#3 showing the responses to 0.3 mM 2-APB, 0.3 mM 2-APB plus 5 mM Mg^2+^ and 3 mM 2-APB. (K) Comparison of relative increase under different conditions. *P* = 4.49E-6 by unpaired student *t*-test. Error bars indicate SEM. **Figure 3 – data source 1** Uncropped, unedited blots for Figure 3B **Figure 3 – data source 2** Uncropped, unedited blots and gels for Figure 3F **Figure 3 – data source 3** Uncropped, unedited blots for Figure 3H **Figure 3 – data source 4** Uncropped, unedited blots for Figure 3I

Alternatively, Mg^2+^ is known as an essential cofactor for enzymatic reactions (de Baaij et al., 2015). Especially, Mg^2+^ is an important regulator of phosphokinases and plays a crucial role in their catalytic activity. Enzymatic/catalytic processes also corroborate the fact that the enhancing effect of Mg^2+^ on TRPV2 took a relatively long time (∼100 s) and could not be immediately eluted (Figure 1A-B). Hence we hypothesize that Mg^2+^ regulates TRPV2 channels through phosphorylation or dephosphorylation. To test this hypothesis, we investigated the phosphorylation level of immunoprecipitated TRPV2 with anti-phosphotyrosine and anti-phospho-Ser/Thr antibody in the presence of 2-APB agonist, with and without Mg^2+^ (Figure 3B). The results revealed a significant increase in tyrosine phosphorylation and serine/threonine phosphorylation levels of TRPV2 in the presence of Mg^2+^. Since the mechanism of phosphorylation involves the transfer of a phosphate (Pi) from ATP to the substrate, we thus used AMP-PNP, a nonhydrolyzable analog of ATP, to replace ATP to inhibit the process of phosphorylation. As shown in Figure 3C, the enhancement effect of Mg^2+^ on TRPV2 currents was abolished when dialyzed AMP-PNP (4 mM) into the cell through recording pipette, suggesting that Mg^2+^ potentiates phosphorylation of TRPV2 upon agonist stimulation.

We next screened the potential kinases involved by treating the cells with various protein kinase inhibitors. As shown in Figure 3D-E, treatment with Ruxolitinib (JAK1 inhibitor) but not MK-2206 (Akt inhibitor), staurosporine (PKC inhibitor), KN93 (CaMKII inhibitor), D4476 (CK1 inhibitor), or U0126 (MEK1/2 inhibitor) abolished the enhancement of TRPV2 activity by Mg^2+^, suggesting that JAK1 is probably the kinase promoting TRPV2 activity.

Utilizing mass spectrometry, we found peptides phosphorylated at the Y335 site that locates on the N terminus (Nt) of TRPV2 (Figure 3 - figure supplement 1). We next tested whether JAK1 directly phosphorylated TRPV2. Based on this finding and considering the difficulty of the purification of the TRPV2 transmembrane region, we purified TRPV2-Nt for *in vitro* phosphorylation experiments. Using *in vitro* kinase assay, we observed that JAK1 directly phosphorylated TRPV2-Nt (Figure 3F).

TRPV2 ion channel has been shown to regulate the phagocytosis of macrophages (Link et al., 2010). We therefore examined macrophage phagocytosis of GFP-expressing *Escherichia coli* (GFP *E. coli*) using flow cytometry by regulating the activity of TRPV2. As expected, blocking TRPV2 by SKF96365 (0.1 mM) significantly inhibited phagocytosis by BMDM cells (74 ± 9% reduction, *n* = 4) (Figure 3G). We then explored whether inhibition of tyrosine phosphorylation by Ruxolitinib affects BMDM phagocytosis. Indeed, Ruxolitinib reduced macrophage phagocytosis in a concentration-dependent manner, with a reduction of 39 ± 2% observed with 10 μM Ruxolitinib (*n* = 3). This result thus corroborates the role of phosphorylation in the functional facilitation of TRPV2 activity.

Next, we evaluated the regulatory effect of JAK1 on TRPV2 function using shRNA-mediated knockdown (Figure 3H). We observed that selective knockdown of JAK1 expression largely reduced Mg^2+^-mediated tyrosine phosphorylation of TRPV2 protein (Figure 3I). Consistently, knockdown of JAK1 expression inhibited the enhancing effect of Mg^2+^ on TRPV2 current responses in BMDM cells (Figure 3J-K). These results together suggest that JAK1 is the kinase underlying Mg^2+^-induced enhancement of TRPV2 activation.

### JAK1 phosphorylates TRPV2 at Y335, Y471, and Y525 molecular sites

Our above results showed that the influx of Mg^2+^ through TRPV2 channel would activate JAK1 and increased the phosphorylation level of the channel, we then investigated the molecular mechanism. Since our mass spectrometry experiment had shown that Y335 was a potential site that may be phosphorylated by JAK1 (Figure 3 - figure supplement 1), we asked whether the mutation at this site would affect the effect of Mg^2+^ on TRPV2 currents. Indeed, mutating Y335 into phenylalanine to simulate dephosphorylation partially inhibited the enhancement of TRPV2 currents by Mg^2+^ (Figure 4A-B). For comparison, the treatment with 5 mM Mg^2+^ increased the 2-APB response (0.3 mM) by approximately 9-fold for mutation Y335F, whereas approximately 16-fold for wild-type TRPV2. The substitution of Y by F approximates a tyrosine that cannot be phosphorylated, while mutations to the negative charge of aspartic acid (D) or glutamic acid (E) are commonly used to mimic phosphorylated tyrosine (Pearlman et al., 2011). As expected, we observed that mutants TRPV2-Y335D and TRPV2-Y335E increased the sensitivity to 2-APB (Figure 4C-D). We thus further verified the effect of Y335F mutation on protein phosphorylation status. Figure 4E illustrates that JAK1-mediated phosphorylation of TRPV2-Nt was abolished by TRPV2(Y335F) and significantly inhibited by the dominant-negative mutant of JAK1 (JAK1-K908A). These data suggest that Y335 is a critical site for JAK1-mediated tyrosine phosphorylation.

**Figure 4.**
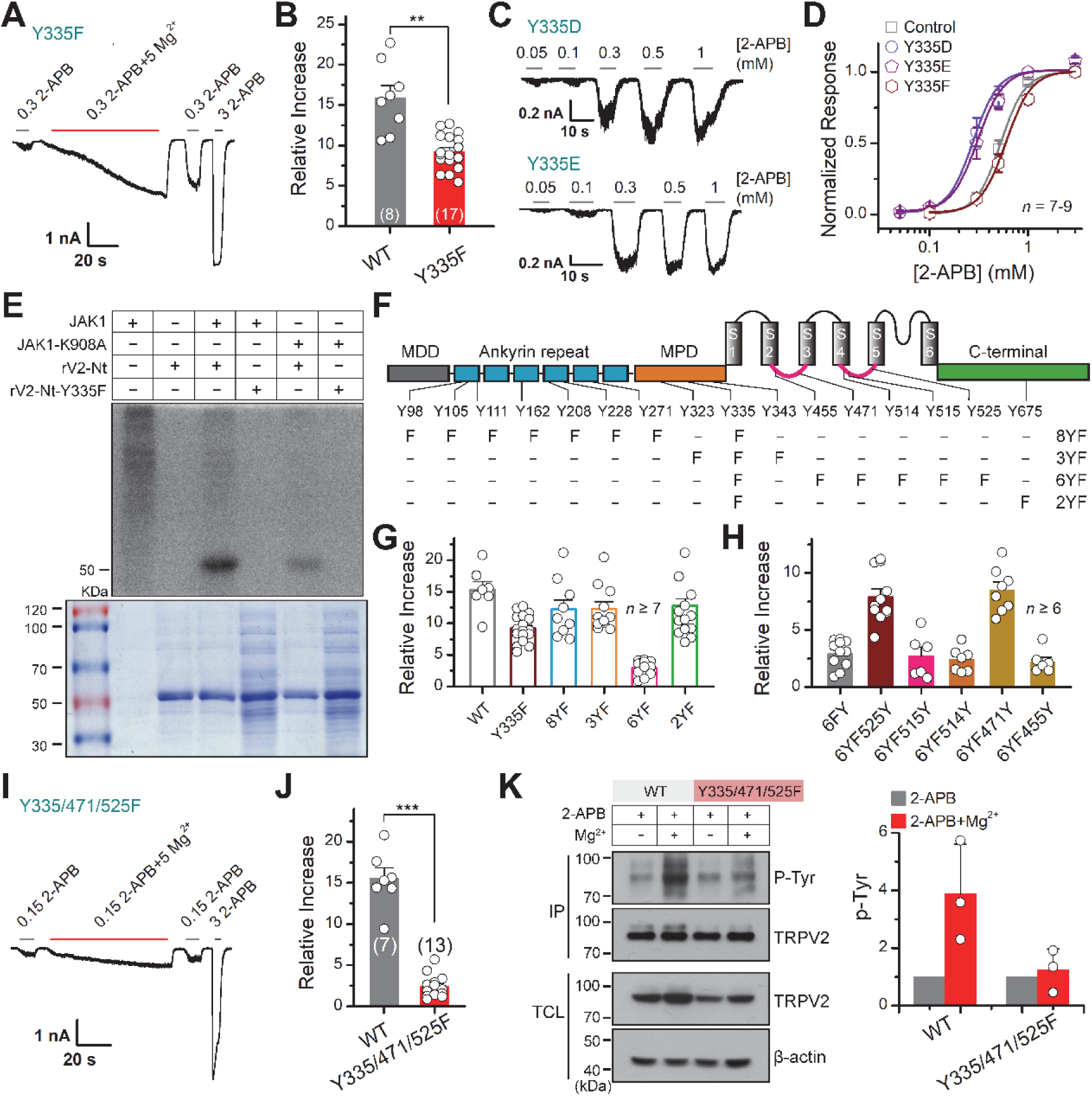
JAK1 has three phosphorylation sites on the TRPV2 channel. (A) Representative whole-cell currents at -60 mV elicited by 0.3 mM 2-APB, 0.3 mM 2-APB plus 5 mM Mg^2+^, and 3 mM 2-APB in HEK293T cells that expressed TRPV2(Y335F). Bars represent duration of stimuli. (B) Comparison of relative changes between wild-type TRPV2 and TRPV2(Y335F) following the treatment by Mg^2+^. *P* = 0.003 by unpaired student *t*-test. (C) Representative whole-cell currents at -60 mV evoked by varying concentrations of 2-APB in HEK293T cells that expressed TRPV2(Y335D) or TRPV2(Y335E). (D) Concentration-response curves of 2-APB for TRPV2 mutants. Solid lines represent fits by a Hill equation with EC_50_ = 0.53 ± 0.01 mM and n_H_ = 3.5 ± 0.1 for TRPV2-WT (*n* = 9); EC_50_ = 0.28 ± 0.01 mM and n_H_ = 3.4 ± 0.2 for Y335D (*n* = 8); EC_50_ = 0.31 ± 0.01 mM and n_H_ = 3.3 ± 0.1 for Y335E (*n* = 7) and EC_50_ = 0.60 ± 0.01 mM and n_H_ = 3.4 ± 0.2 for Y335F (*n* = 8). (E) *In vitro* kinase assay with [^32^P]-γ-ATP, immunoprecipitated tyrosine kinase JAK1 and recombinant His-tagged wild-type or mutant TRPV2 N-terminus. Phosphorylation signals were examined by autoradiography. (F) Linear diagram of the TRPV2 channel topology, with all intracellular tyrosine residues labeled, and a summary of substitutions of tyrosine by phenylalanine used in this study. (G) Summary plot of the Mg^2+^-dependent enhancement in various mutants. All the TRPV2 mutants retained their normal responses to 2-APB. (H) Statistic results for the Mg^2+^-dependent enhancement for mutants which were respectively reverse mutated from TRPV2-6YF. (I) Representative whole-cell currents at -60 mV elicited by 0.15 mM 2-APB, 0.15 mM 2-APB plus 5 mM Mg^2+^, and 3 mM 2-APB in HEK293T cells that expressed TRPV2-Y335/471/525F. (J) Average plot of the relative changes of wild-type and Y335/471/525F currents following treatment by Mg^2+^. *P* = 2.30E-9 0.001 by unpaired student *t*-test. (K) Immunoblotting analysis with anti-phosphotyrosine antibody (pTyr) showing the tyrosine phosphorylation levels in HEK 293T cells transfected with TRPV2 or TRPV2-Y335/471/525F in the absence and presence of Mg^2+^. *Right*, quantitative analysis of the fold increase of tyrosine-phosphorylated TRPV2 proteins and TRPV2(Y335/471/525F) proteins following different treatments (*n* = 3; means ± S.D.). Error bars indicate SEM. **Figure 4 – data source 1** Uncropped, unedited blots and gels for Figure 4E **Figure 4 – data source 2** Uncropped, unedited blots for Figure 4K

Since mutation Y335F partially abolishes the enhancement effect of Mg^2+^, there may exist other phosphorylation sites in TRPV2 channel protein. Using mutant Y335F as a template, we further mutated the tyrosine residues in the N-terminal ankyrin repeat domain (ARD), the membrane-proximal domain (MPD), intracellular linkers (Linker), and the C-terminal (Ct) into phenylalanine by site-directed mutagenesis, respectively. We obtained the following mutants: 8YF (Y98/105/111/162/208/228/271/335F), 3YF (Y323/335/343F), 6YF (Y335/455/471/514/515/525F), and 2YF (Y335/675F) (Figure 4F). Mutant 6YF greatly reduced the Mg^2+^ induced enhancement of TRPV2 response (Figure 4G). When phenylalanine at positions 471 and 525 were reversed back to tyrosine from the 6YF mutant (6YF471Y and 6YF525Y), the enhancement of TRPV2 was rescued (Figure 4H).

Triple mutant TRPV2(Y335/471/525F) was generated to confirm the significance of these three specific sites. The results in Figure 4I-J displayed that TRPV2(Y335/471/525F) largely eliminated the enhancement of TRPV2 by Mg^2+^. The protein sequence alignment showed that Y335, Y471, and Y525 amino acid residues are highly conserved in various mammalian TRPV2 homologs (Figure 4 – figure supplement 1). Moreover, this tri-mutant also downregulated tyrosine phosphorylation levels of immunoprecipitated TRPV2 protein (Figure 4K).

### Tyrosine phosphorylation enhances chemical and thermal sensitization of TRPV2

Protein phosphorylation is a reversible post-translational modification mediated by kinases and phosphatases. Having characterized JAK1 as the kinase for tyrosine phosphorylation of TRPV2, we next sought to identify the phosphatases that counteracted this process. We took advantage of various protein phosphatase inhibitors to search for the phosphatases that mediated the dephosphorylation of TRPV2. The protein phosphatases comprise the phosphoprotein phosphatase (PPP) family, the protein phosphatase Mg^2+^- or Mn^2+^-dependent (PPM) family, and the protein tyrosine phosphatase (PTP) (Barford et al., 1998). We first examined the effect of pretreatment of the phosphatase inhibitors, which would elevate the basal phosphorylation level of TRPV2 and compromise the subsequent enhancing effect of Mg^2+^ on current responses. As shown in Figure 5A-B, a significant impact was observed with PTP inhibitor 1 (2-bromo-4’-hydroxy acetophenone) and PTP inhibitor 2 (2-bromo-1-(4-methoxyphenyl)-ethanone), but not PPP inhibitors salubrinal, LB-100, cyclosporin A, cantharidin, nor the PPM inhibitor CCT007093. We then confirmed that inhibition of tyrosine dephosphorylation by PTP inhibitors indeed increased tyrosine phosphorylation levels of TRPV2 (Figure 5C-D). Besides, we found that in BMDM, the upregulation of tyrosine phosphorylation of TRPV2 caused by PTP inhibitors induced a left-shift of the concentration-response curve to agonist application (Figure 5E-F). The corresponding EC_50_ values were 0.18 ± 0.01 mM and 0.09 ± 0.01 mM in the presence of PTP inhibitor 1 or 2, respectively, compared to EC_50_ = 0.55 ± 0.01 mM under control condition. Conversely, TRPV2(Y335/471/525F) mutant deficit in Mg^2+^ influx showed no significant change in the presence of PTP inhibitors (Figure 5G).

**Figure 5.**
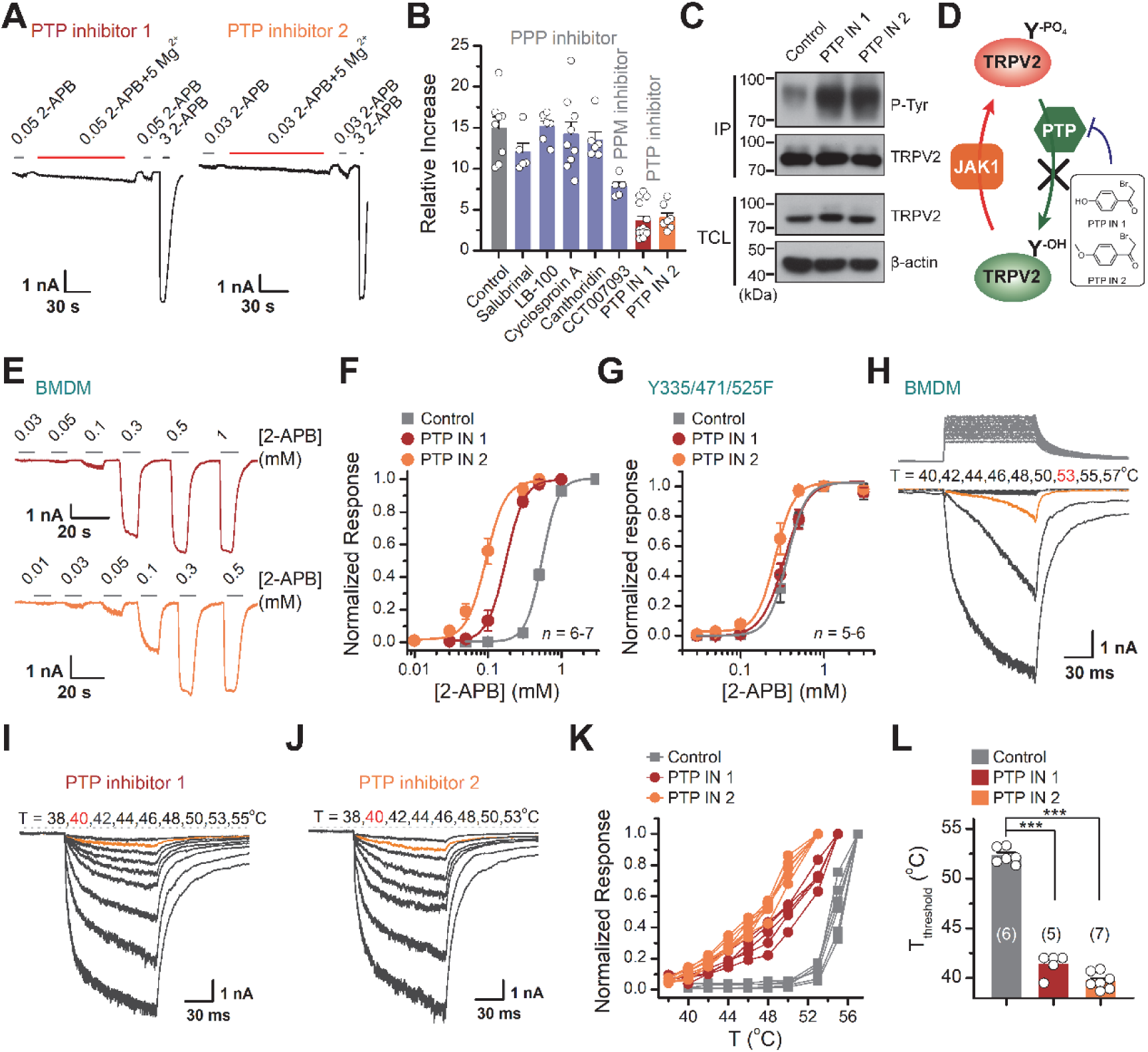
Increasing the phosphorylation level of TRPV2 by inhibition of dephosphorylase activity enhances the channel sensitivity to its stimuli. (A) Whole-cell recordings from TRPV2-expressing HEK293T cell were consecutively challenged with 0.3 mM 2-APB, 0.3 mM 2-APB plus 5 mM Mg^2+^ and 3 mM 2-APB. The cells were pretreated with protein tyrosine phosphatase (PTP) inhibitor 1 and PTP inhibitor 2 for 5 min, respectively. (B) Summary plot of effects of various phosphatase inhibitors on TRPV2 currents. (C) Immunoblotting analysis with anti-phosphotyrosine antibody exhibiting tyrosine phosphorylation of immunoprecipitated TRPV2-Flag in HEK293T cells under control conditions and after treatment with PTP inhibitor 1 or PTP inhibitor 2. (D) Schematic diagram showing increased TRPV2 tyrosine-phosphorylation levels caused by phosphokinase JAK1 or inhibition of PTP activity. (E) Representative whole-cell currents evoked by increasing concentrations of 2-APB for rBMDMs. The cells were pre-treated with PTP inhibitor 1 (*Top*) and PTP inhibitor 2 (*Bottom*). (F) Dose-response curves of 2-APB. Fitting by Hill’s equation resulted in the following: EC_50_ = 0.55 ± 0.01 mM and n_H_ = 3.9 ± 0.2 for control (*n* = 6); EC_50_ = 0.18 ± 0.01 mM and n_H_ = 3.4 ± 0.1 for treatment by PTP inhibitor 1 (*n* = 6) and EC_50_ = 0.09 ± 0.01 mM and n_H_ = 3.3 ± 0.3 for treatment by PTP inhibitor 2 (*n* = 7). (G) Concentration-response curves of 2-APB in TRPV2-Y335/471/525F-expressing HEK 293T cells under treatment by DMSO, PTP inhibitor 1 or PTP inhibitor 2. Fitting by Hill’s equation resulted in the following: EC_50_ = 0.36 ± 0.01 mM and n_H_ = 3.8 ± 0.1 for control (*n* = 5); EC_50_ = 0.34 ± 0.01 mM and n_H_ = 3.1 ± 0.1 for treatment by PTP inhibitor 1 (*n* = 6) and EC_50_ = 0.26 ± 0.01 mM and n_H_ = 3.8 ± 0.7 for treatment by PTP inhibitor 2 (*n* = 6). (H-L) Representative responses to a family of rapid temperature jumps for rBMDMs under control (H), and inhibition by PTP inhibitor 1 (I) or PTP inhibitor 2 (J). (K) Temperature-dependent response curves, measured from the maximal currents at the end of temperature steps. Each cure indicates measurements from an individual cell. (L) Comparison of temperature threshold (*T*_threshold_). *T*_threshold_ = 52.3 ± 0.3°C for control (*n* = 6), *T*_threshold_ = 40.8 ± 0.7°C for treatment by PTP inhibitor 1 (*n* = 5) and *T*_threshold_ = 39.7 ± 0.3 °C for treatment by PTP inhibitor 2 (*n* = 7). *P* = 1.49E-11 for *T*_threshold_ of control vs. PTP inhibitor 1 treatment and *P* = 1.19E-12 for *T*_threshold_ of control vs. PTP inhibitor 2 treatment using one-way ANOVA t-test. Error bars indicate SEM. **Figure 5 – data source 1** Uncropped, unedited blots for Figure 5C

We next determined the effect of PTP-mediated dephosphorylation of TRPV2 on its temperature sensitivity. We employed an ultrafast infrared laser system capable of delivering a short temperature pulse surrounding BMDMs. Figure 5H-J illustrates representative heat-activated currents of TRPV2 treated with DMSO (Figure 5H), PTP inhibitor 1 (Figure 5I), and PTP inhibitor 2, respectively (Figure 5J). The current-temperature relationship in Figure 5K confirms that the inhibition of dephosphorylase activity caused a significantly left-shifted temperature dependence curve and displayed a much shallower slope. Remarkably, we observed that boosting tyrosine phosphorylation lowered by ∼12 °C the thermal activation threshold of TRPV2 (Figure 5L). Similar results were obtained for TRPV2 channels expressed in HEK 293T heterologous expression systems (Figure 5 - figure supplement 1). Taken together, these results support that tyrosine phosphorylation promotes both the chemical and thermal sensitivities of TRPV2, which are both controlled by phosphatase dephosphorylation.

### PTPN1 phosphatase controls tyrosine phosphorylation homeostasis

We further determined the subtypes of PTP phosphatases involved in controlling TRPV2 phosphorylation processes. We observed that knocking down of PTPN1 phosphatase by shRNA increased the tyrosine phosphorylation of TRPV2 (Figure 6A-B), which increased its sensitivity to the chemical agonist 2-APB (Figure 6C). Conversely, no effect was observed following the inhibition of the expression of PTPN2, PTPN11, PTPN12, PTPN14, PTP4A1, or PTEN (Figure 6C). As corroboration, downregulating PTPN1 expression to boost the basal phosphorylation level compromised the enhancing effect of subsequently applied Mg^2+^ on TRPV2 current responses (Figure 6D-E).

**Figure 6.**
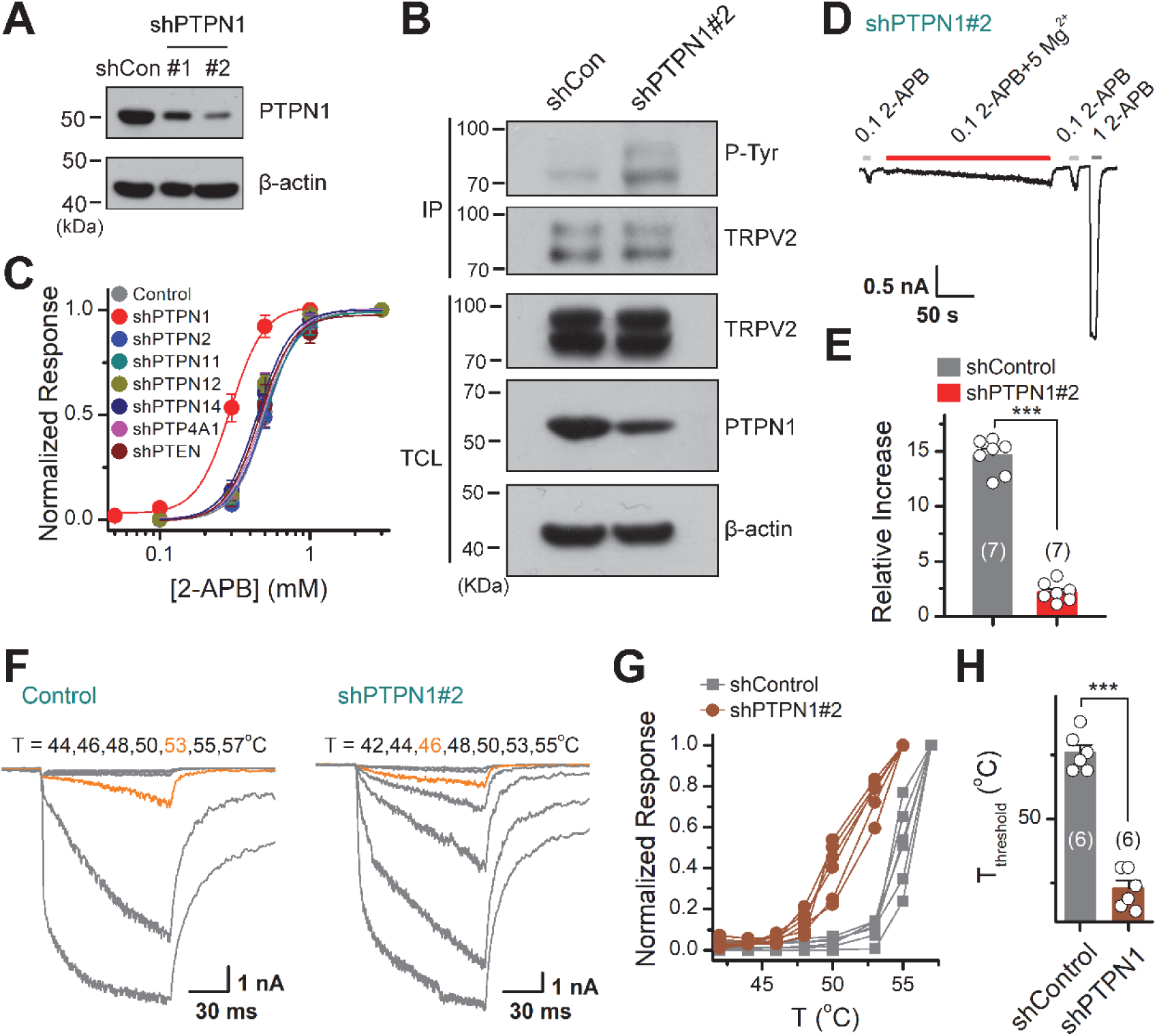
PTPN1 is a phosphatase that mediates the dephosphorylation of TRPV2. (A) Immunoblot analysis (with anti-PTPN1 or anti-β-action) of HEK 293T cells transfected for 48 h with PTPN1-targeting shRNA (shPTPN1#1 and shPTPN1#2) or shCon to test knockdown efficiency of shRNA. (B) Immunoblot analysis of the tyrosine phosphorylation level of TRPV2 in HEK293T cells transfected with shControl or shPTPN1#2 for 48 h. (C) Concentration-response curves of 2-APB. Whole-cell recordings were performed in HEK 293T transfected with various protein tyrosine phosphatase-targeting shRNA. (D) Whole-cell recordings in TRPV2-expressing HEK293T cells that transfected for 48 h with shPTPN1#2 showing the response to 0.1 mM 2-APB, 0.1 mM 2-APB plus 5 mM Mg^2+^ and 1 mM 2-APB. (E) Comparison of relative changes under different conditions. *P* = 3.88E-10 by unpaired student *t*-test. (F) Effects of inhibition of PTPN1 on temperature dependence. Representative responses to a family of rapid temperature jumps in TRPV2-expressing HEK 293T transfected for 48 h with shControl (*left*) or shPTPN1#2 (*right*). (G) Comparison of current-temperature relationships. Temperature response curves were measured from the maximal currents at the end of each temperature step and each curve indicates measurements from an individual cell. (H) Comparison of temperature threshold (*T*_threshold_). *T*_threshold_ = 52.6 ± 0.3 °C (*n* = 6) for transfected with shConrtrol; *T*_threshold_ = 47.3 ± 0.3 °C (*n* = 6) for transfected with shPTPN1#2. *P* = 1.99E-7 by unpaired student *t*-test. Error bars represent SEM. **Figure 6 – data source 1** Uncropped, unedited blots for Figure 6A **Figure 6 – data source 2** Uncropped, unedited blots for Figure 6B

We then investigated the effect of PTPN1 on heat activation of TRPV2, by applying time-locked temperature jumps. Increasing tyrosine phosphorylation by inhibition of the PTPN1-mediated dephosphorylation significantly decreased the temperature threshold of TRPV2 activation (Figure 6F-H). These data suggest that PTPN1 phosphatase restrains basal phosphorylation levels of TRPV2 to regulate its function.

## Discussion

TRPV2 ion channel senses a wide range of sensory inputs and is an essential player in physiopathological contexts. In the present study, we delineate a hitherto unrecognized tyrosine phosphorylation module that defines the homeostatic sensitivity of TRPV2 ion channel (Figure 6 – figure supplement 1).

Our data show that Mg^2+^ modulates tyrosine phosphorylation levels of the TRPV2 channel protein thereby its current responses. This observation mirrors the established role of Mg^2+^ in the regulation of phosphokinase catalytic activities and the regulation of diverse ion channels including NMDA receptors (Antonov and Johnson, 1999) and TRP ion channels (Cao et al., 2014; Lee et al., 2005; Luo et al., 2012; Obukhov and Nowycky, 2005; Yang et al., 2014). We reveal that Mg^2+^-mediated enhancing effect on TRPV2 current responses is tuned by JAK1 kinase and PTPN1 phosphatase at Y335, Y471, and Y525 molecular sites. Tyrosine phosphorylation of TRPV2 controls not only its sensitivity to chemical stimulations, but also its thermal activation threshold. Temperature sensing is essential to survive and adapt since failure to avoid noxious temperatures can cause fundamental tissue damage. TRPV1, TRPV2, TRPV3, TRPV4, and TRPM2 channels together sense a broad temperature range spanning from physiological warmness to noxious hotness. The physiological role of the TRPV1 channels in thermosensation has been demonstrated by the knock-out of the TRPV1 channels in mice (Garami et al., 2011). However, the physiological role of the TRPV2 channels remains unclear while it is responsive to noxious heat (>52 °C) in heterologous systems. We here demonstrate that enhancing the tyrosine phosphorylation levels of TRPV2 protein lowers its thermal threshold to a near-body temperature level (∼40 °C). TRPV2 might act as a heat thermosensor in physio-pathological conditions when encountering either or both Mg^2+^ surges and upregulated tyrosine phosphorylation (Yu et al., 2011). For instance, intracellular free Mg^2+^ can be increased by adenosine triphosphate (ATP) depletion induced by either mitochondrial deficits (Kubota et al., 2003) or cell reactive states that consume a high amount of cytosolic ATP (Brocard et al., 1993; Gaussin et al., 1997). In addition to tyrosine phosphorylation, oxidation of methionine residues or other potential endogenous modulators would independently or synergistically modulate TRPV2 channel sensitivity (Fricke et al., 2019).

Protein post-translational modification represents a main endogenous regulatory mechanism of ion channels and immune signaling, by changing the plasma membrane expression or altering the biophysical properties of the channels. PKA-mediated phosphorylation of the TRPV1 channels and the TRPV2 channels have been proposed (Jeske et al., 2008; Stokes et al., 2004). Phosphorylation of TRPV1 channels via PKC-related pathway or Src-related pathway was reported to mediate TRPV1 surface expression level (Studer and McNaughton, 2010; Zhang et al., 2005). Differentially, our data suggest that tyrosine phosphorylation of TRPV2 directly alters its biophysical properties without changing the expression of TRPV2 on the plasma membrane.

Mg^2+^ participates in a wide range of fundamental cellular reactions and its deficiency may lead to many disorders. It has been reported that magnesium deficiency caused by deficiency genetic deficiencies in *MAGT1* impairs anti-virus immune response which can be restored by intracellular free magnesium supplementation (Chaigne-Delalande et al., 2013). This study also shows that the concentration of intracellular free Mg^2+^ can be increased by long-term Mg^2+^ supplementation. As a more efficient way to alter intracellular Mg^2+^ concentrations, Mg^2+^ can permeate into the cell through ion channels such as TRPM6, TRPM7, or/and magnesium transports like MagT1 (Deason-Towne et al., 2011; Goytain and Quamme, 2005; Voets et al., 2004). Using TRPV2 mutant deficient in Mg^2+^ permeation and patch clamp glass pipette-guided Mg^2+^-chelator EDTA supplying, our data suggest that transient Mg^2+^ buildup on the intracellular side is required for shifting the tyrosine phosphorylation level. This mechanism differs from the action of Mg^2+^ on TRPV1 channels, where a high concentration of Mg^2+^ potentiates the TRPV1 activity from the extracellular side but inhibits TRPV1 currents from the intracellular side (Cao et al., 2014; Yang et al., 2014).

By specifically perturbing the JAK1-mediated phosphorylation and PTPN1-mediated dephosphorylation, we could substantially alter the chemical and thermal sensitivity of TRPV2 ion channel. Thus, TRPV2 channel sensitivity is maintained at the homeostatic point by dynamically balanced phosphorylation/dephosphorylation processes. The Mg^2+^-enhanced TRPV2 current responses are quickly reverted (Figure 1A-F), suggesting that the endogenous phosphatase activity of PTPN1 is high. As such, TRPV2 is likely maintained at a low level of phosphorylation in basal conditions.

## Materials and methods

### Key resources table

2 Key resources table
| Reagent type (species) or resource | Designation | Source or reference | Identifiers | Additional information |
| --- | --- | --- | --- | --- |
| Antibody | Rabbit anti-Phosphotyrosine antibody | Abcam | Cat#ab179530 | WB (1:1000) |
| Antibody | Rabbit anti-Phospho-(Ser/Thr)Phe antibody | Abcam | Cat#ab17464; RRID: AB_443891 | WB (1:1000) |
| Antibody | Rabbit anti-Flag antibody | Proteintech | Cat#20543-1-AP; RRID: AB_11232216 | WB (1:3000) |
| Antibody | Rabbit anti-TRPV2 antibody | Alomone Labs | Cat#ACC-032; RRID: AB_2040266 | WB (1:500), IP (1:200) |
| Antibody | Goat anti-mouse IgG (H+L) | Jackson ImmunoResearch | Cat#115-035-003; RRID: AB_10015289 |  |
| Antibody | Goat anti-rabbit IgG (H+L) | Jackson ImmunoResearch | Cat#111-005-003; RRID: AB_2337913 |  |
| Antibody | Rabbit anti-JAK1 antibody | Abcam | Cat#ab133666 | WB (1:1000) |
| Antibody | Rabbit anti-PTPN1 antibody | Abcam | Cat#ab244207; RRID: AB_2877148 | WB (1:1000) |
| Antibody | *ProteinIso*<sup>®</sup> Protein G Resin | TransGen | Cat#DP401 |  |
| Antibody | Mouse anti-Flag Affinity Gel | Bimake | Cat#B23102; RRID: AB_2728745 |  |
| Chemical compound, drug | 2-APB | Sigma-Aldrich | Cat#:D9754, CAS: 524-95-8 | TRPV2 agonist |
| Chemical compound, drug | MgCl<sub>2</sub>·6H<sub>2</sub>O | Sigma-Aldrich | Cat#:M2393, CAS: 7791-18-6 |  |
| Chemical compound, drug | Na<sub>2</sub>-ATP | Sigma-Aldrich | Cat#A2383; CAS: 34369-07-8 |  |
| Chemical compound, drug | EDTA | Biosharp | Cat#BS107; CAS: 60-00-4 |  |
| Chemical compound, drug | AMP-PNP | Sigma-Aldrich | Cat#A2647;<br>CAS: 25612-73-1 |  |
| Chemical compound, drug | MK-2206 | TargetMol | Cat#T1952;<br>CAS: 1032350-13-2 | Akt inhibitor |
| Chemical compound, drug | Staurosporine | TargetMol | Cat#T6680;<br>CAS: 62996-74-1 | PKC inhibitor |
| Chemical compound, drug | KN-93 Phosphate | TargetMol | Cat#T2606;<br>CAS: 1188890-41-6 | CaMKII inhibitor |
| Chemical compound, drug | D4476 | TargetMol | Cat#T2449;<br>CAS: 301836-43-1 | CK1 inhibitor |
| Chemical compound, drug | U0126-EtOH | TargetMol | Cat#T6223;<br>CAS: 1173097-76-1 | MEK1/2 inhibitor |
| Chemical compound, drug | Ruxolitinib | TargetMol | Cat#T1829;<br>CAS: 941678-49-5 | JAK1 inhibitor |
| Chemical compound, drug | Salubrinal | TargetMol | Cat#T3045;<br>CAS: 405060-95-9 | PP1 inhibitor |
| Chemical compound, drug | LB-100 | MCE | Cat#HY-18597;<br>CAS: 1632032-53-1 | PP2A inhibitor |
| Chemical compound, drug | Cyclosporin A | TargetMol | Cat#T0945;<br>CAS: 59865-13-3 | PP2B inhibitor |
| Chemical compound, drug | Cantharidin | Aladdin | Cat#c111020;<br>CAS: 56-25-7 | PP1 and PP2A inhibitor |
| Chemical compound, drug | CCT007093 | TargetMol | Cat#T1927;<br>CAS:176957-55-4 | PPM1D inhibitor |
| Chemical compound, drug | PTP inhibitor 1 | TargetMol | Cat#T7084;<br>CAS: 2491-38-5 | PTPs inhibitor |
| Chemical compound, drug | PTP inhibitor 2 | TargetMol | Cat#T7541;<br>CAS: 2632-13-5 | PTPs inhibitor |
| Cell lines | HEK293T | ATCC | Cat#CRL-3216;<br>RRID: CVCL_0063 |  |
| Software and algorithms | QStudio |  |  | Developed by Dr. Feng Qin from University of New York at Buffalo |
| Software and algorithms | Micro-Manager 1.4 | Vale lab, UCSF |  |  |
| Software and algorithms | Clampfit | Molecular Devices, Sunnyvale, CA |  |  |
| Software and algorithms | IGOR | Wavemetrics, Lake Oswego, OR, USA |  |  |
| Software and algorithms | SigmaPlot | SPSS Science, Chicago, IL |  |  |
| Software and algorithms | OriginPro | OriginLab Corporation, MA, USA |  |  |
| Software and algorithms | ImageJ | Schneider et al., 2012 |  |  |

### Cell lines

HEK 293T cells, CHO cells, Hela cells, and ND7/23 cells were grown in Dulbecco’s modified Eagle’s medium (DMEM, Thermo Fisher Scientific, MA) containing 4.5 mg/ml glucose, 10% heat-inactivated fetal bovine serum (FBS), 1% penicillin-streptomycin, and were incubated at 37°C in a 5% CO_2_ humidified incubator. Cells grown into ∼80% confluence were transfected with the desired DNA constructs using lipofectamine 2000 (Invitrogen, Carlsbad, CA) following the protocol provided by the manufacturer. Transfected cells were reseeded on poly-L-lysine coated glass coverslips for electrophysiological experiments. Experiments took place usually 12–24 h after transfection.

### cDNA constructs and mutagenesis

WT mouse TRPV2 (mTRPV2), rat TRPV2 (rTRPV2) were generously provided by Dr. Feng Qin (State University of New York at Buffalo, Buffalo, USA). JAK1 was a gift from Dr. Hongbing Shu (Medical Research Institute, Wuhan University). All mutations were generated using the overlap-extension polymerase chain reaction method as previously described (Wang et al., 2020) and were verified by DNA sequencing. Oligo DNAs targeting JAK1, PTPN1, and several PTPs were synthesized, annealed, and inserted into pLKO.1 vector. The sequences of JAK1 shRNA are as follows: for rat JAK1 shRNA: #1, 5′-GCCCTGAGTTACTTGGAAGAT -3′; #2, 5′-CGGTCCAATCTGCACAGAATA -3′; #3, 5′-GCAGAAACCAAATGTTCTTCC-3′; for human JAK1 shRNA: #1, 5′-GAGACTTCCATGTTACTGATT -3′; #2, 5′-GACAGTCACAAGACTTGTGAA -3′; #3, 5′-GCCTTAAGGAATATCTTCCAA-3′. The sequences of PTPN1 shRNA are as follows: for human PTPN1 shRNA: #1, 5′-TGCGACAGCTAGAATTGGAAA-3′; #2, 5′-GCTGCTCTGCTATATGCCTTA-3′.

### Rat and mouse bone marrow-derived macrophages

Bone marrow-derived cells were isolated from 4-8 weeks old Sprague-Dawley (SD) rats as described (Zhang et al., 2020). After the rats were euthanized, the femurs and tibias were collected. The cells were resuspended in bone marrow differentiation media, RPMI1640 supplemented with 1% penicillin-streptomycin, 10% FBS, and 30% L929 cells conditioned medium containing macrophage colony stimulating factor (M-CSF) for 4-6 d to obtain BMDMs. Cells were cultured at 37 °C in a classic CO_2_ incubator with 5% CO_2_.

All animals were housed in the specific pathogen-free animal facility at Wuhan University and all animal experiments were following protocols approved by the Institutional Animal Care and Use Committee of Wuhan University (NO. WDSKY0201804) and adhered to the Chinese National Laboratory Animal-Guideline for Ethical Review of Animal Welfare. The animals were euthanatized with CO_2_ followed by various studies.

### Preparation of dorsal root ganglion (DRG) neurons

DRG neurons were prepared for electrophysiological experiments by minor modification of a previously described method (Tian et al., 2019). Briefly, 4-6 week-old adult SD male rats were deeply anesthetized and decapitated. DRGs together with dorsal-ventral roots and attached spinal nerves were isolated from thoracic and lumbar segments of spinal cords. After removal of the attached nerves and surrounding connective tissues, DRG neurons were rinsed with ice-cold phosphate buffer saline (PBS). Ganglia were dissociated by enzymatic treatment with collagenase type IA (1 mg/ml), trypsin (0.4 mg/ml) and DNase I (0.1 mg/ml) and incubated at 37 °C for 30 min. Then cells were dispersed by gentle titration, collected by centrifuge, seeded onto 0.1 mg ml^−1^ poly-L-lysine–coated coverslips, maintained in DMEM/F12 medium containing 10% FBS, 1% penicillin, and streptomycin. Electrophysiology recordings were carried out ∼2–4 h after plating.

### Electrophysiology

The patch-clamp recording of channel currents was made in either whole-cell or inside-out configuration. Currents were amplified using an Axopatch 200B amplifier (Molecular Devices, Sunnyvale, CA) through a BNC-2090/MIO acquisition system (National Instruments, Austin, TX). Data acquisition was controlled by QStudio developed by Dr. Feng Qin at State University of New York at Buffalo. Data were typically sampled at 5 kHz and low-pass filtered at 1 kHz. Recording pipettes were pulled from borosilicate glass capillaries (World Precision Instruments, WPI) to 2–4 MΩ when filled with 150 mM NaCl solution. The compensation of pipette series resistance (> 80%) and capacitance was taken by using the built-in circuitry of the amplifier, and the liquid junction potential between the pipette and bath solutions was zeroed prior to seal formation. All voltages were defined as membrane potentials with respect to extracellular solutions. For whole-cell recording, the bath solution contained the following (in mM) 140 NaCl, 5 KCl, 3 EGTA, 10 HEPES (the pH was adjusted to 7.4 with NaOH). In one set of experiments, the salt of YCl_2_ (Y means Mg^2+^, Mn^2+^, Ca^2+^, Ba^2+^, Zn^2+^, Cu^2+^, Ni^2+^, Cd^2+^ or Co^2+^) was individually dissolved in deionized water to make stock solutions and subsequently diluted into a basic solution ([in mM] 140 NaCl, 5 KCl and 10 HEPES, pH 7.4) to make a desired final concentration. The solution containing 10-100 mM Mg^2+^ was prepared from 140 mM NaCl-containing solution by replacing the appropriate NaCl with MgCl_2_. The internal pipette solution consisted of (in mM): 140 CsCl, 10 HEPES, and 1 ATP-Na_2_, pH 7.4 (adjusted with CsOH). For inside-out recordings, the bath and pipette solutions were symmetrical and contained (in mM) 140 NaCl, 5 KCl, 10 HEPES, pH 7.4 adjusted with NaOH. Channel activators were diluted into the recording solution at the desired final concentrations and applied to the cell of interest through a gravity-driven local perfusion system. Unless otherwise stated, all chemicals were purchased from Sigma (Sigma, St. Louis, MO). Water-insoluble reagents were dissolved in either 100% ethanol or DMSO to make stock solutions and were diluted in the recording solutions at appropriate concentrations before experiments. The final concentrations of ethanol or DMSO did not exceed 0.3%, which did not affect the currents. All experiments except those for heat activation were sampled at room temperature (22–24 °C).

### Temperature jump

Fast-temperature jumps were produced by a single emitter infrared laser diode (1470 nm) as previously described (Yao et al., 2009). Briefly, the laser diode was driven by a pulsed quasi-CW current power supply (Stone Laser, Beijing, China), and the pulsing of the controller was controlled from a computer through the data acquisition card using QStudio software. Constant temperature steps were generated by irradiating the tip of an open pipette filled with the pipette solution and the current of the electrode was used as a readout for feedback control. The sequence of the modulation pulses was stored and subsequently played back to apply temperature jumps to the cell of interest. The temperature was calibrated off-line from the pipette current based on the temperature dependence of electrolyte conductivity. The threshold temperature for heat activation of TRPV2 was determined as the temperature at which ∼10% of its maximum response was induced.

### Ca^2+^ imaging

Fluorescent images of HEK 293T cells co-expressed with GCaMP6m (a gift from Dr. Liangyi Chen, Peking Unversity) together with TRPV2-WT or TRPV2-E609/614Q were acquired under an inverted epifluorescence microscope (Olympus IX 73, Tokyo, Japan) equipped with a complete illumination system (Lambda XL, Sutter Instruments). Intracellular Ca^2+^ was measured using a cool CCD camera (CoolSNAP ES2, Teledyne Photometrics) which was controlled by Micro-Manager 1.4 (Vale lab, UCSF) at 470 ± 22 nm excitation. The extracellular solution contained with 140 mM NaCl, 5 mM KCl, 1.8 mM CaCl_2_, and 10 mM HEPES, pH 7.4. Changes in intracellular Ca^2+^ levels were calculated by subtracting the basal fluorescence intensity (mean value collected for 10 s before agonist addition) from the fluorescence intensity after exposure to agonist.

### Immunoprecipitation and Western blot

In brief, cells were collected and lysed in Nonidet P-40 lysis buffer containing 150 mM NaCl, 1 mM EDTA, 1% Nonidet P-40, 1% protease inhibitor cocktail, and 1% phosphatase inhibitor cocktail if needed after washing with PBS. The anti-Flag affinity gel or the appropriate antibodies were added into the lysates and incubated at 4 °C for 4 h or overnight with slow rotation. After being washed three times with prelysis buffer containing 500 mM NaCl, the precipitants were resuspended into 2× SDS sample buffer, boiled, subjected to SDS-polyacrylamide gel electrophoresis (SDS-PAGE). Immunoblot analysis was performed with the appropriate antibodies.

### Mass Spectrometry Analysis

To identify *in vivo* tyrosine phosphorylation sites of TRPV2, HEK 293T cells were transfected with Flag-tagged TRPV2. After 24 hours, the cells were harvested following the treatment with 0.3 mM 2-APB or the combination of 0.3 mM 2-APB and 5 mM Mg^2+^ lasting for 5 min. Flag-TRPV2 was immunoprecipitated by anti-Flag affinity gel and subjected to SDS-PAGE.

The samples were analyzed by liquid chromatography–tandem mass spectrometry (LC-MS/MS) using a Q Exactive-HF mass spectrometer (Thermo Fisher Scientific). The LC-MS/MS data were processed using Proteome Discoverer (Thermo Fisher Scientific) and searched against the Swiss-prot Homo sapiens protein sequence database. Data were analyzed using ProteinPilot software (AB SCIEX).

### In vitro kinase assay

*In vitro* kinase assay was performed as previously described (Li et al., 2019). In brief, HEK 293T cells were transfected with plasmids encoding Flag-JAK1, Flag-JAK1(K908A), respectively. Cells were lysed with NP-40 lysis buffer and the cell lysates were immunoprecipitated with anti-Flag agarose (Sigma, St. Louis, MO). His-tagged TRPV2 and His-tagged TRPV2 (Y335F) were purified from bacteria (*E. coli*) using Ni-Agarose Resin. For the JAK1 *in vitro* kinase assay in Figure 3, Flag-JAK1 was respectively incubated with His-TRPV2 in the kinase buffer (6.25 mM Tris-HCl [pH7.5], 0.125 mM Na_3_VO_4_, 2.5 mM MgCl_2_, 0.125 mM EGTA, 0.625 mM DTT, and 0.01% Triton X-100) in the presence of 10 μCi [^32^P]-γ-ATP (Perkin Elmer Company) with a final volume of 20 μl. For the JAK1 *in vitro* kinase assay in Figure 4, His-TRPV2 and His-TRPV2 (Y335F) were incubated with or without Flag-JAK1 and Flag-JAK1(K908A) in the kinase buffer in the presence of 10 μCi [^32^P]-γ-ATP with a final volume of 20 μl. The mixture was incubated at 30 °C on a shaker with 300 rpm shaking for 60 min. The reaction mixtures were resolved by SDS-PAGE, and ^32^P-labelled proteins were analyzed by autoradiography.

### Assessment of phagocytosis

For phagocytosis assays, BMDMs were incubated with RPMI 1640 medium addition of E.coli-GFP together with 0.1 or 0.05 mM SKF96365, or 2, 5, and 10 μM Ruxolitinib in 6-well translucent plates (JET Biofil, China) for 2 h at 37 °C. After washing 2-3 times by PBS, the BMDMs were harvested by cell Scrapers, resuspended into PBS, and analyzed by flow cytometry using a CytoFLEX Flow Cytometer (Beckman Coulter, USA).

### Statistical analysis

Electrophysiological data were analyzed offline with Clampfit (Molecular Devices, Sunnyvale, CA), IGOR (Wavemetrics, Lake Oswego, OR, USA), SigmaPlot (SPSS Science, Chicago, IL, USA), and OriginPro (OriginLab Corporation, MA, USA). For concentration dependence analysis, the modified Hill equation was used: Y = A1 + (A2 - A1) / [1 + 10^(logEC_50_ – X)*n_H_], in which EC_50_ is the half-maximal effective concentration, and n_H_ is the Hill coefficient. All data are expressed as either mean ± standard error (SEM) or mean ± standard (SD) as stated, from a population of cells (*n*). Statistical tests of significance were carried out by Student’s *t*-test for one-group comparison and two-group comparison or one-way analysis of variance (ANOVA) tests for multiple group comparisons, and *P* < 0.05 was considered statistically significant (\**P* < 0.05, ** *P* < 0.01, *** *P* < 0.001).

## Data availability

All major datasets supporting the conclusions of this article has been deposited at Dryad, https://doi.org/10.5061/dryad.41ns1rng6 (Jing Yao et al., 2022).

## Acknowledgements

We are grateful to Drs. Xiaolu Zhao, Zan Huang, Yan Wang and members of Yao lab for critical comments and helpful discussions. We also would like to thank the core facilities of College of Life Sciences at Wuhan University for technical help. This work was supported by grants from the National Natural Science Foundation of China (31929003, 31830031, 32171147, 31871174 and 31671209), and the Fundamental

Research Funds for the Central Universities (2042021KF0218).

## Author contributions

J.Y. designed and supervised the study. X.M., P.P., Y.W., D.J., M.Z., Y.L., P.W., Q.G., and J.Y. carried out the experiments and analyzed data. C.X., H-N.D., B.Z. and D.L. provided technical support and suggestions. X.M., P.P., and J.Y. wrote the paper with inputs from all other authors. The authors read and approved the final manuscript.

---

**Figure 1 - figure supplement 1.**
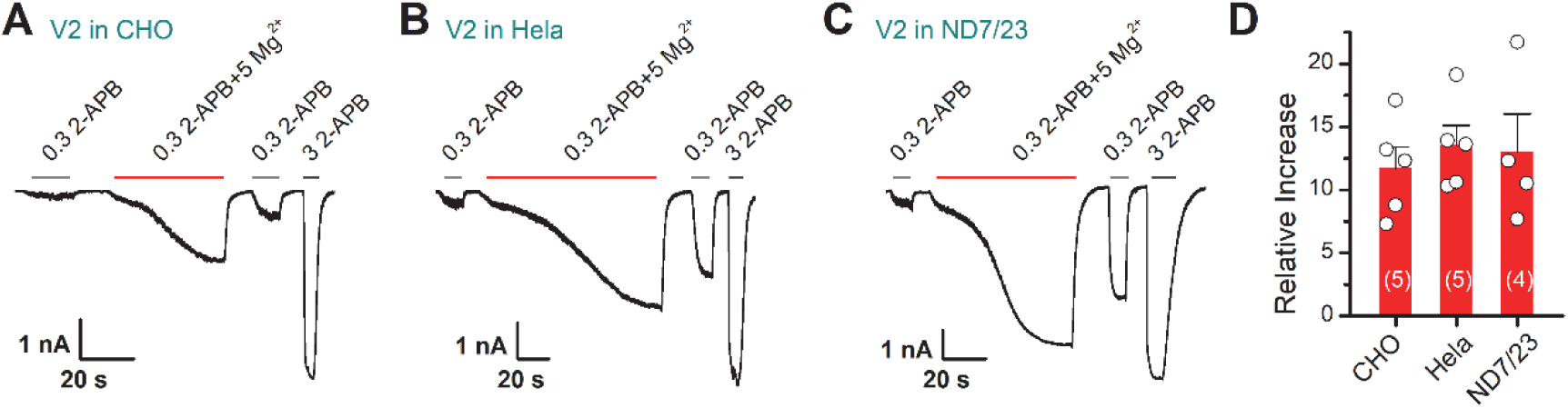
Mg^2+^ potentiates TRPV2 currents expressed in various cell lines. (A-C) Representative whole-cell recordings in CHO, Hela, or ND7/23 that expressed TRPV2 showing the response to 0.3 mM 2-APB, 0.3 mM 2-APB plus 5 mM Mg^2+^ and 3 mM 2-APB. (D) Summary of relative currents induced by 0.3 mM 2-APB and 0.3 mM 2-APB plus 5 mM Mg^2+^.

**Figure 1 - figure supplement 2.**
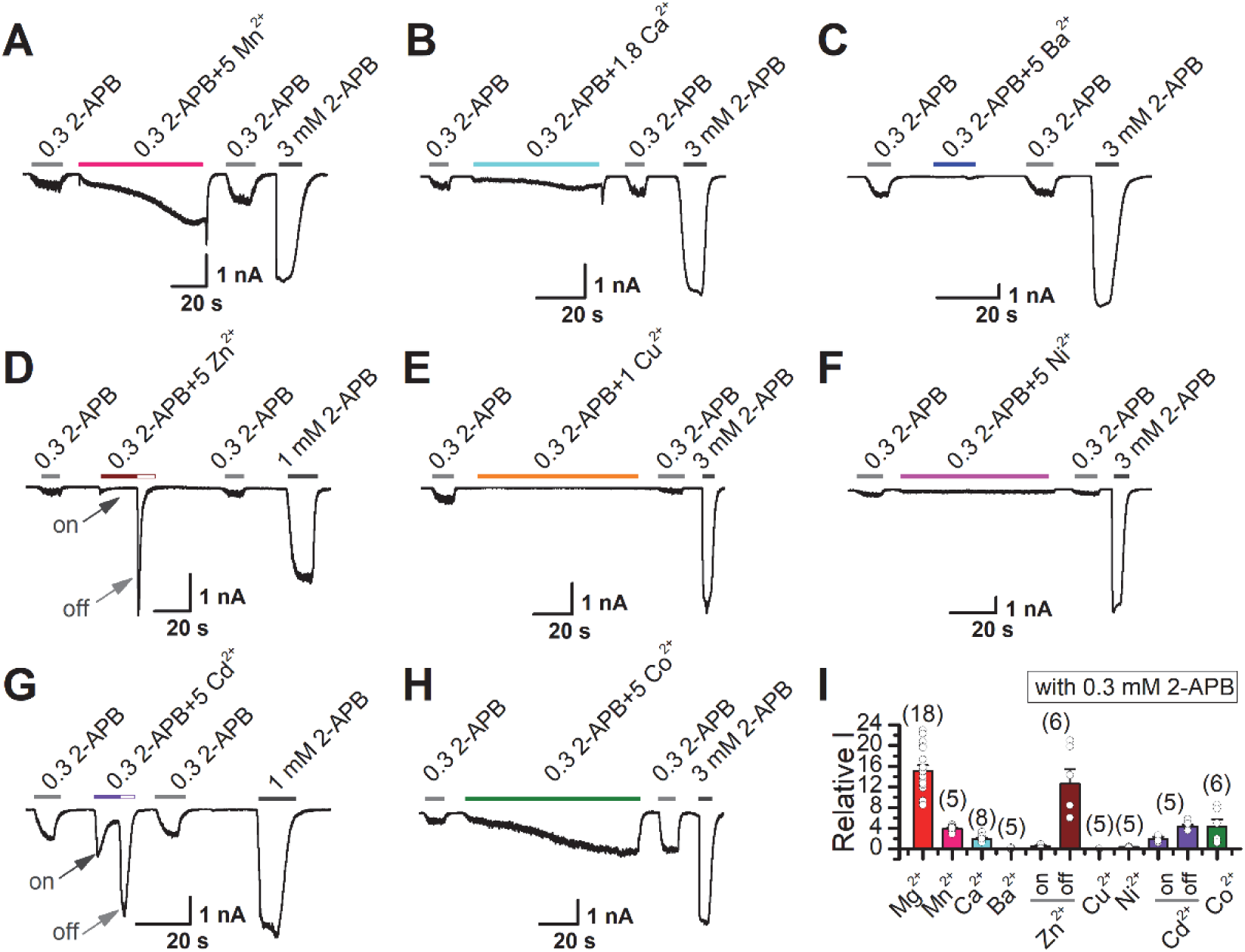
Effects of various divalent cations on 2-APB-evoked TRPV2 currents. (A-H) Representative whole-cell currents in TRPV2-expressing HEK 293T cells induced by 0.3 mM 2-APB, the combination of 0.3 mM 2-APB and various divalent cations Mn^2+^ (A), Ca^2+^ (B), Ba^2+^ (C), Zn^2+^ (D), Cu^2+^ (E), Ni^2+^ (F), Cd^2+^ (G), and Co^2+^ (H), respectively. (I) Summary of relative currents evoked by the combination of 0.3 mM 2-APB and different divalent cations versus 0.3 mM 2-APB only.

**Figure 2 - figure supplement 1.**
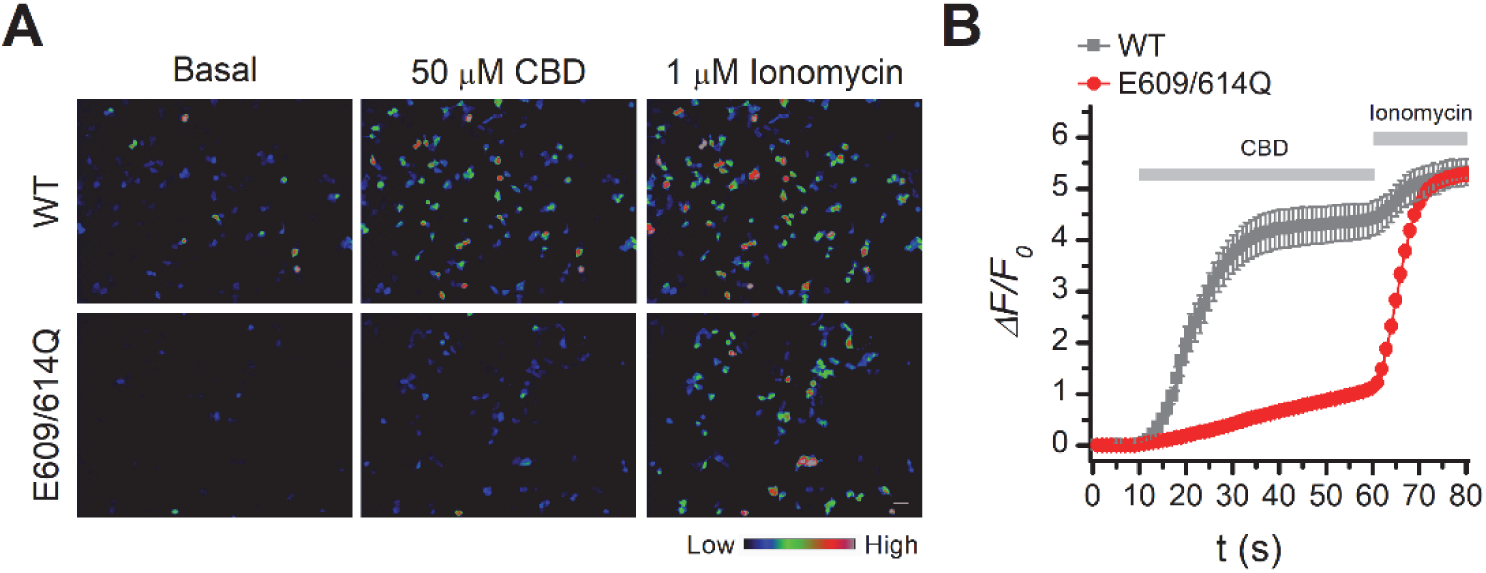
Ca^2+^ imaging in TRPV2(WT) or TRPV2(E609/614Q) expressing HEK 293T cells. (A) Ca^2+^ responses of TRPV2(WT) (upper) or TRPV2(E609/614Q) (lower) expressing HEK 293T cells were following exposure to 50 μM cannabidiol (CBD) and 1 μM ionomycin. Scale bar, 50 μm. (B) Averaged responses of TRPV2(WT) (gray, *n* = 28) or TRPV2(E609/614Q) (red, *n* = 58) transfected cells exposed to CBD and ionomycin. GCaMP fluorescence changes were computed as (F_i_–F_0_)/F_0_, where F_i_ represented fluorescence intensity at any frame and F_0_ was the baseline fluorescence calculated from the averaged fluorescence of the first 10 s.

**Figure 3 - figure supplement 1.**
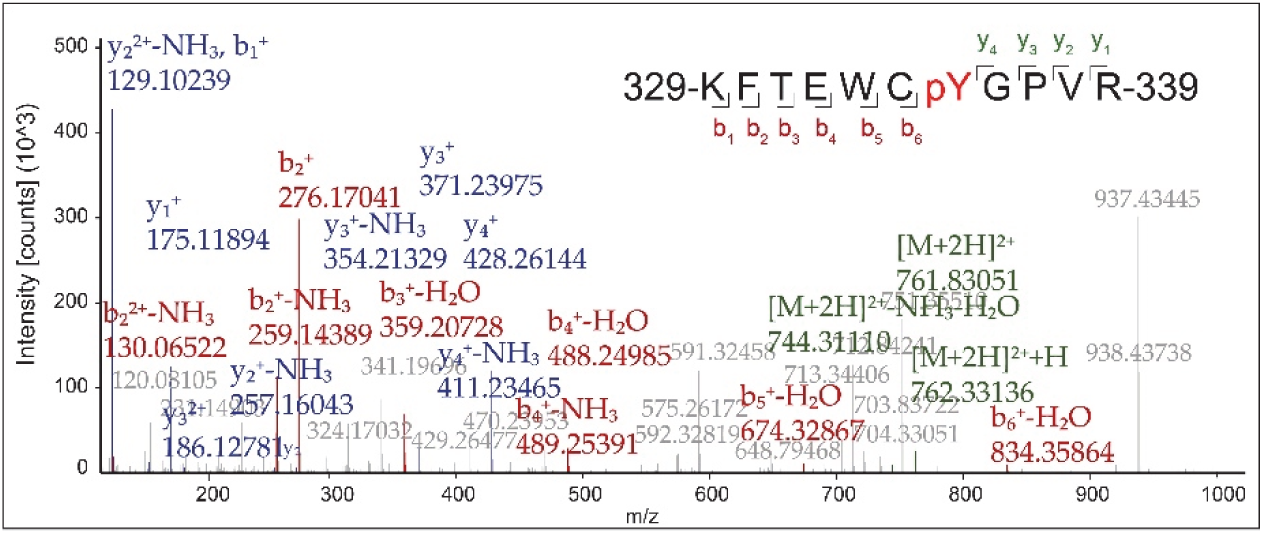
Mass spectrometry analysis of the phosphorylation of TRPV2. Mass spectrometry analysis showing the phosphorylation of TRPV2 in HEK 293T cells after treatment by 0.3 mM 2-APB plus 5 mM Mg^2+^, followed by immunoprecipitation (with anti-FLAG agarose). MS/MS ion spectrum with the matched b and y ions of the pY335-containing tryptic peptide KFTEWCpYGPVR was shown.

**Figure 4 - figure supplement 1.**
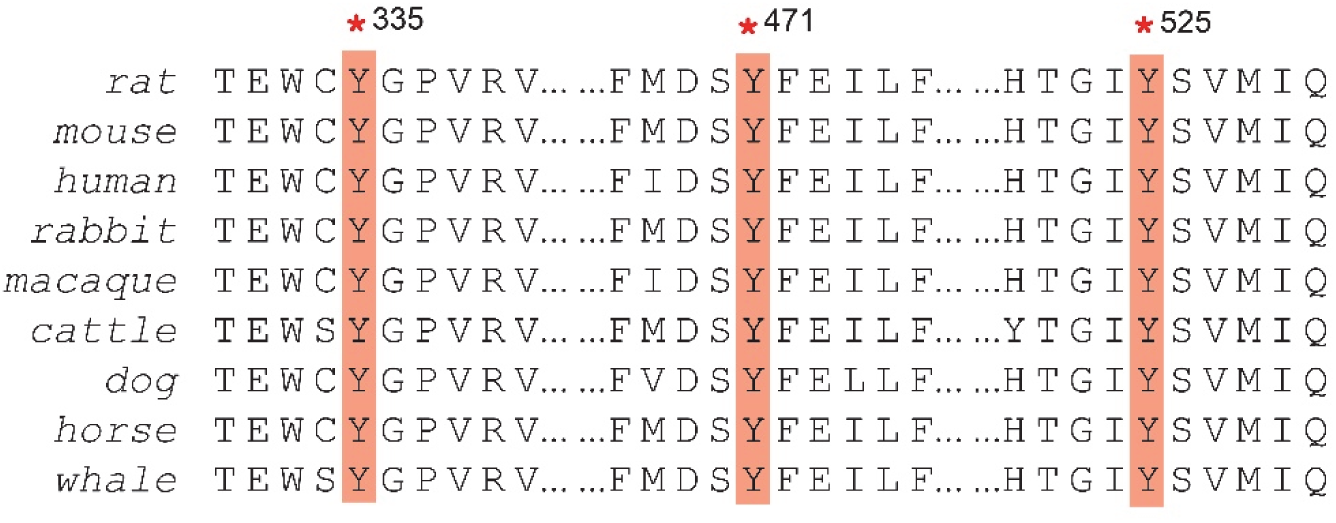
Partial amino acid sequence alignment of TRPV2 channels. Multiple alignments of TRPV2 amino acid sequences surrounding Y335, Y471, and Y525 from rat, mouse, human, rabbit, macaque, cattle, dog, horse, and whale. The residues of Y335, Y471, and Y525 are boxed in the sequence alignment.

**Figure 5 - figure supplement 1.**
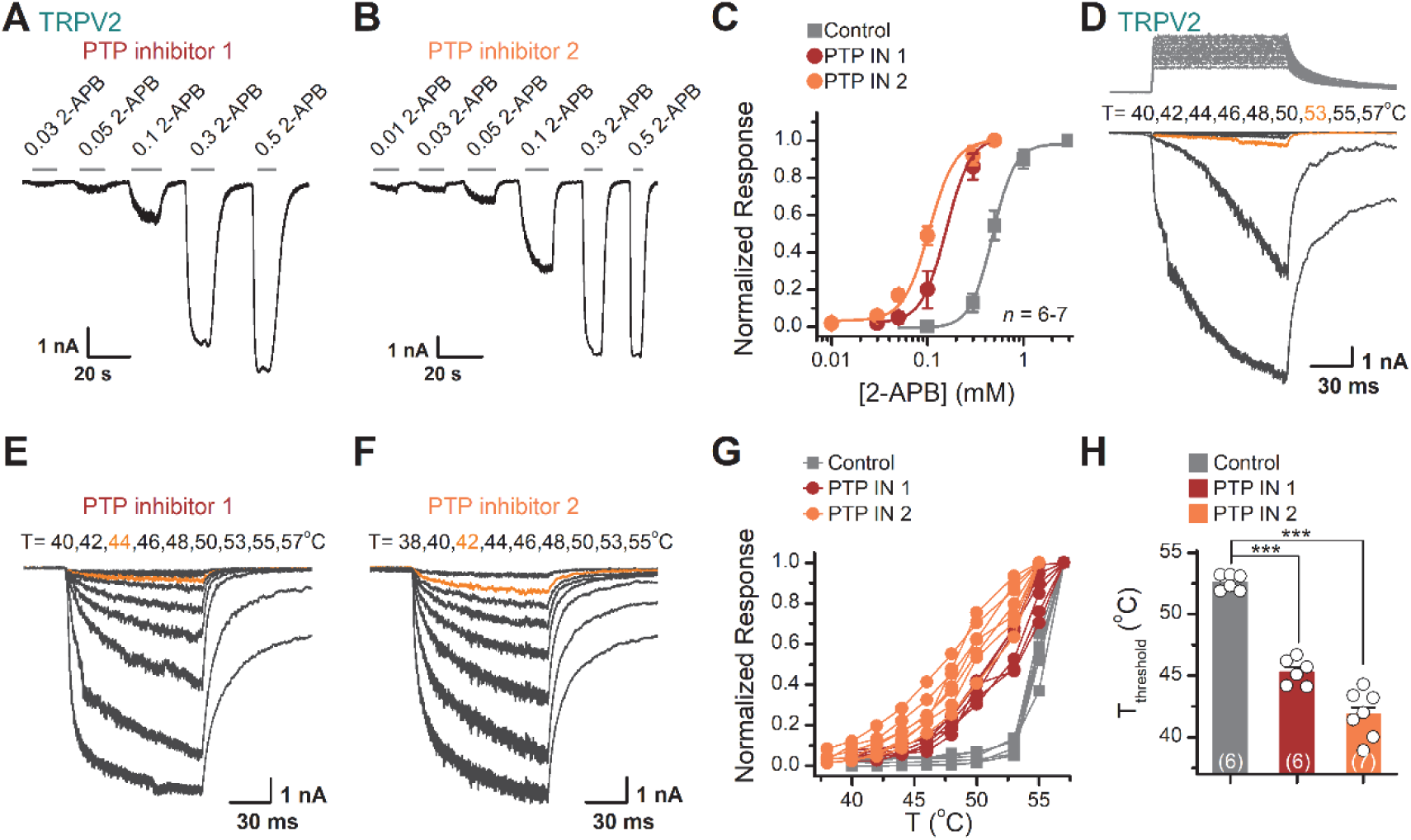
Inhibition of PTP activity by inhibitors enhanced the TRPV2 sensitivity to 2-APB and heat in TRPV2-expressing HEK 293T cells. (A-B) Representative whole-cell recordings from TRPV2-expressing HEK 293T cells pre-treated with PTP inhibitor 1 (A) and PTP inhibitor 2 (B). The cells were exposed to increasing concentrations of 2-APB and the holding potential was -60 mV. (C) Concentration-response curves of 2-APB. Solid lines indicate fits by a Hill’s equation, with EC_50_ = 0.48 ± 0.02 mM and n_H_ = 3.7 ± 0.4 for control (*n* = 7); EC_50_ = 0.16 ± 0.01 mM and n_H_ = 3.4 ± 0.2 for the treatment by PTP inhibitor 1 (*n* = 6), and EC_50_ = 0.10 ± 0.01 mM and n_H_ = 3.2 ± 0.3 for the treatment by PTP inhibitor 2 (*n* = 7). (D-F) Representative whole-cell currents evoked by a family of rapid temperature jumps under control condition (D), the treatment by PTP inhibitor 1 (E) or PTP inhibitor 2 (F). (G) Temperature-dependent response curves, measured from the maximal currents at the end of temperature steps. Each cure represents measurements from an individual cell. (H) Comparison of T_threshold_. *T_threshold_* = 52.6 ± 0.3 °C for control condition (*n* = 6), *T_threshold_* = 45.3 ± 0.4 °C for the treatment by PTP inhibitor 1 (*n* = 6) and *T_threshold_* = 41.9 ± 0.7 °C for the treatment by PTP inhibitor 2 (*n* = 7). *P* = 9.21E-8 for *T*_threshold_ of control vs. PTP inhibitor 1 treatment and *P* = 2.11E-10 for *T*_threshold_ of control vs. PTP inhibitor 2 treatment using one-way ANOVA t-test. Error bars indicate SEM.

**Figure 6 - figure supplement 1.**
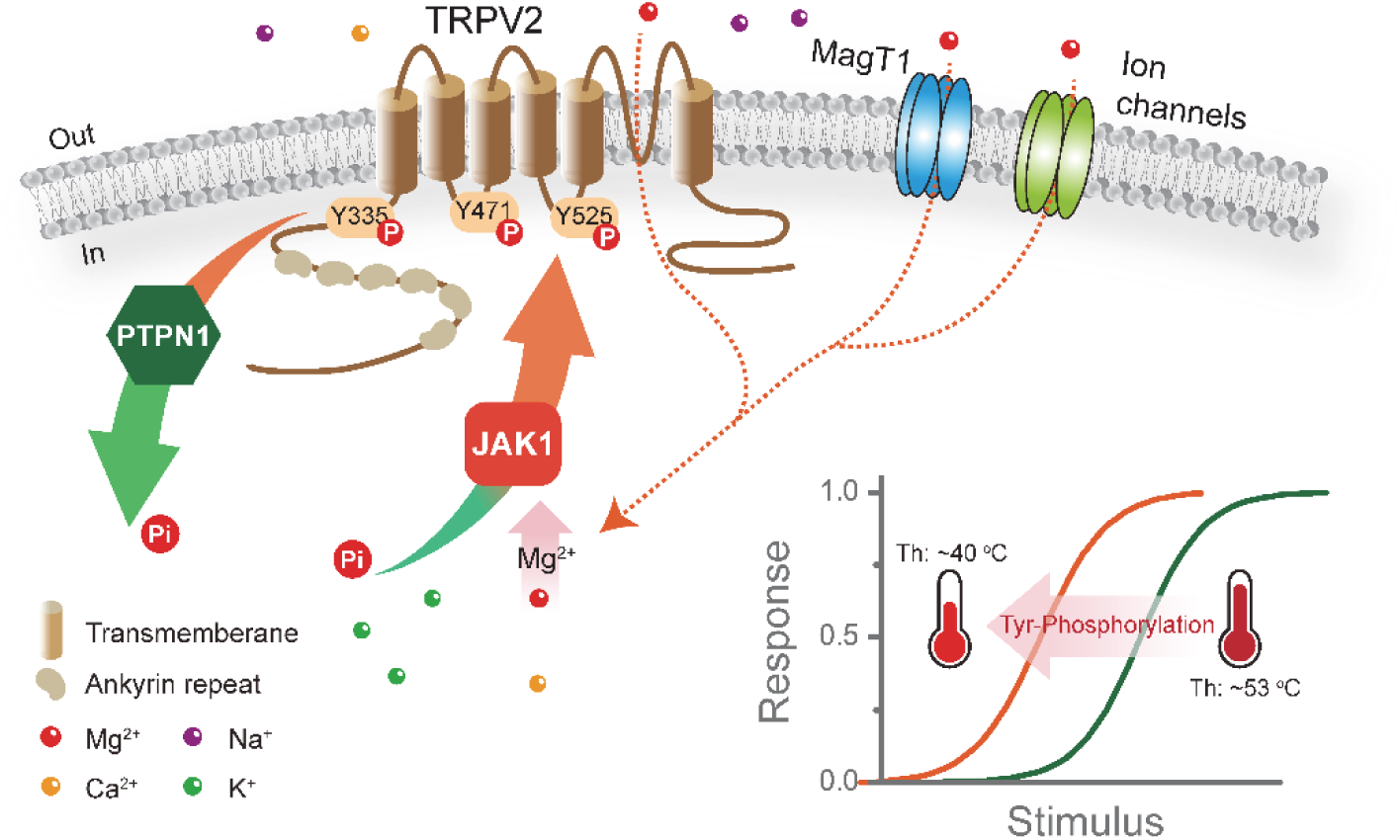
Tyrosine phosphorylation sets the agonist and heat sensitivity of TRPV2. This study demonstrates that JAK1 phosphokinase mediates Mg^2+^-dependent phosphorylation of TRPV2 at Y335, Y471, and Y525 residues. And, increasing tyrosine phosphorylation of TRPV2 lowers its thermal activation threshold and enhances its sensitivity to agonistic stimuli. Furthermore, PTPN1 is the tyrosine phosphatase that mediates the dephosphorylation of the TRPV2 channel.

## References

1. Antonov, S.M., and Johnson, J.W. (1999). Permeant ion regulation of N-methyl-D-aspartate receptor channel block by Mg^2+^. Proc Natl Acad Sci U S A 96: 14571–14576.

2. Aoyagi, K., Ohara-Imaizumi, M., Nishiwaki, C., Nakamichi, Y., and Nagamatsu, S. (2010). Insulin/phosphoinositide 3-kinase pathway accelerates the glucose-induced first-phase insulin secretion through TrpV2 recruitment in pancreatic beta-cells. Biochem J 432: 375–386.

3. Bang, S., Kim, K.Y., Yoo, S., Lee, S.H., and Hwang, S.W. (2007). Transient receptor potential V2 expressed in sensory neurons is activated by probenecid. Neurosci Lett 425: 120–125.

4. Barford, D., Das, A.K., and Egloff, M.P. (1998). The structure and mechanism of protein phosphatases: insights into catalysis and regulation. Annu Rev Biophys Biomol Struct 27: 133–164.

5. Brocard, J.B., Rajdev, S., and Reynolds, I.J. (1993). Glutamate-induced increases in intracellular free Mg^2+^ in cultured cortical neurons. Neuron 11: 751–757.

6. Cao, X., Ma, L., Yang, F., Wang, K., and Zheng, J. (2014). Divalent cations potentiate TRPV1 channel by lowering the heat activation threshold. J Gen Physiol 143: 75–90.

7. Caterina, M.J., Rosen, T.A., Tominaga, M., Brake, A.J., and Julius, D. (1999). A capsaicin-receptor homologue with a high threshold for noxious heat. Nature 398: 436–441.

8. Chaigne-Delalande, B., Li, F.Y., O’Connor, G.M., Lukacs, M.J., Jiang, P., Zheng, L., Shatzer, A., Biancalana, M., Pittaluga, S., Matthews, H.F., et al. (2013). Mg^2+^ regulates cytotoxic functions of NK and CD8 T cells in chronic EBV infection through NKG2D. Science 341: 186–191.

9. de Baaij, J.H., Hoenderop, J.G., and Bindels, R.J. (2015). Magnesium in man: implications for health and disease. Physiol Rev 95: 1–46.

10. De Petrocellis, L., Ligresti, A., Moriello, A.S., Allarà, M., Bisogno, T., Petrosino, S., Stott, C.G., and Di Marzo, V. (2011). Effects of cannabinoids and cannabinoid-enriched Cannabis extracts on TRP channels and endocannabinoid metabolic enzymes. Br J Pharmacol 163: 1479–1494.

11. Deason-Towne, F., Perraud, A.L., and Schmitz, C. (2011). The Mg^2+^ transporter MagT1 partially rescues cell growth and Mg^2+^ uptake in cells lacking the channel-kinase TRPM7. FEBS Lett 585: 2275–2278.

12. Entin-Meer, M., Cohen, L., Hertzberg-Bigelman, E., Levy, R., Ben-Shoshan, J., and Keren, G. (2017). TRPV2 knockout mice demonstrate an improved cardiac performance following myocardial infarction due to attenuated activity of peri-infarct macrophages. PLoS One 12: e0177132.

13. Fricke, T.C., Echtermeyer, F., Zielke, J., de la Roche, J., Filipovic, M.R., Claverol, S., Herzog, C., Tominaga, M., Pumroy, R.A., Moiseenkova-Bell, V.Y., et al. (2019). Oxidation of methionine residues activates the high-threshold heat-sensitive ion channel TRPV2. Proceedings of the National Academy of Sciences 116: 24359–24365.

14. Garami, A., Pakai, E., Oliveira, D.L., Steiner, A.A., Wanner, S.P., Almeida, M.C., Lesnikov, V.A., Gavva, N.R., and Romanovsky, A.A. (2011). Thermoregulatory phenotype of the Trpv1 knockout mouse: thermoeffector dysbalance with hyperkinesis. J Neurosci 31: 1721–1733.

15. Gaussin, V., Gailly, P., Gillis, J.M., and Hue, L. (1997). Fructose-induced increase in intracellular free Mg^2+^ ion concentration in rat hepatocytes: relation with the enzymes of glycogen metabolism. Biochem J 326 ( *Pt 3*): 823–827.

16. Goytain, A., and Quamme, G.A. (2005). Identification and characterization of a novel mammalian Mg^2+^ transporter with channel-like properties. BMC Genomics 6: 48.

17. Hisanaga, E., Nagasawa, M., Ueki, K., Kulkarni, R.N., Mori, M., and Kojima, I. (2009). Regulation of calcium-permeable TRPV2 channel by insulin in pancreatic beta-cells. Diabetes 58: 174–184.

18. Hu, H.Z., Gu, Q., Wang, C., Colton, C.K., Tang, J., Kinoshita-Kawada, M., Lee, L.Y., Wood, J.D., and Zhu, M.X. (2004). 2-aminoethoxydiphenyl borate is a common activator of TRPV1, TRPV2, and TRPV3. J Biol Chem 279: 35741–35748.

19. Hu, J., Gao, Y., Huang, Q., Wang, Y., Mo, X., Wang, P., Zhang, Y., Xie, C., Li, D., and Yao, J. (2021). Flotillin-1 Interacts With and Sustains the Surface Levels of TRPV2 Channel. Front Cell Dev Biol 9: 634160.

20. Huynh, K.W., Cohen, M.R., Jiang, J., Samanta, A., Lodowski, D.T., Zhou, Z.H., and Moiseenkova-Bell, V.Y. (2016). Structure of the full-length TRPV2 channel by cryo-EM. Nat Commun 7: 11130.

21. Iwata, Y., Ito, S., Wakabayashi, S., and Kitakaze, M. (2020). TRPV2 channel as a possible drug target for the treatment of heart failure. Lab Invest 100: 207–217.

22. Jeske, N.A., Diogenes, A., Ruparel, N.B., Fehrenbacher, J.C., Henry, M., Akopian, A.N., and Hargreaves, K.M. (2008). A-kinase anchoring protein mediates TRPV1 thermal hyperalgesia through PKA phosphorylation of TRPV1. Pain 138: 604–616.

23. Jing Yao, Xiaoyi Mo, Peiyuan Pang, Yulin Wang, Dexiang Jiang, Mengyu Zhang, Yang Li, Peiyu Wang, Qizhi Geng, Chang Xie, et al. Tyrosine phosphorylation tunes chemical and thermal sensitivity of TRPV2 ion channel Dryad, https://doi.org/10.5061/dryad.41ns1rng6 (2022).

24. Juvin, V., Penna, A., Chemin, J., Lin, Y.L., and Rassendren, F.A. (2007). Pharmacological characterization and molecular determinants of the activation of transient receptor potential V2 channel orthologs by 2-aminoethoxydiphenyl borate. Mol Pharmacol 72: 1258–1268.

25. Kanzaki, M., Zhang, Y.Q., Mashima, H., Li, L., Shibata, H., and Kojima, I. (1999). Translocation of a calcium-permeable cation channel induced by insulin-like growth factor-I. Nature Cell Biology 1: 165–170.

26. Katanosaka, Y., Iwasaki, K., Ujihara, Y., Takatsu, S., Nishitsuji, K., Kanagawa, M., Sudo, A., Toda, T., Katanosaka, K., Mohri, S., et al. (2014). TRPV2 is critical for the maintenance of cardiac structure and function in mice. Nat Commun 5: 3932.

27. Kubota, T., Tokuno, K., Nakagawa, J., Kitamura, Y., Ogawa, H., Suzuki, Y., Suzuki, K., and Oka, K. (2003). Na^+^/Mg^2+^ transporter acts as a Mg^2+^ buffering mechanism in PC12 cells. Biochemical and Biophysical Research Communications 303: 332–336.

28. Lee, J., Cha, S.K., Sun, T.J., and Huang, C.L. (2005). PIP2 activates TRPV5 and releases its inhibition by intracellular Mg^2+^. J Gen Physiol 126: 439–451.

29. Li, W., Wang, H.Y., Zhao, X., Duan, H., Cheng, B., Liu, Y., Zhao, M., Shu, W., Mei, Y., Wen, Z., et al. (2019). A methylation-phosphorylation switch determines Plk1 kinase activity and function in DNA damage repair. Sci Adv 5: eaau7566.

30. Link, T.M., Park, U., Vonakis, B.M., Raben, D.M., Soloski, M.J., and Caterina, M.J. (2010). TRPV2 has a pivotal role in macrophage particle binding and phagocytosis. Nat Immunol 11: 232–239.

31. Liu, B., and Qin, F. (2016). Use Dependence of Heat Sensitivity of Vanilloid Receptor TRPV2. Biophys J 110: 1523–1537.

32. Luo, J., Stewart, R., Berdeaux, R., and Hu, H. (2012). Tonic inhibition of TRPV3 by Mg^2+^ in mouse epidermal keratinocytes. J Invest Dermatol 132: 2158–2165.

33. McGahon, M.K., Fernandez, J.A., Dash, D.P., McKee, J., Simpson, D.A., Zholos, A.V., McGeown, J.G., and Curtis, T.M. (2016). TRPV2 Channels Contribute to Stretch-Activated Cation Currents and Myogenic Constriction in Retinal Arterioles. Invest Ophthalmol Vis Sci 57: 5637–5647.

34. Muraki, K., Iwata, Y., Katanosaka, Y., Ito, T., Ohya, S., Shigekawa, M., and Imaizumi, Y. (2003). TRPV2 is a component of osmotically sensitive cation channels in murine aortic myocytes. Circ Res 93: 829–838.

35. Nagasawa, M., Nakagawa, Y., Tanaka, S., and Kojima, I. (2007). Chemotactic peptide fMetLeuPhe induces translocation of the TRPV2 channel in macrophages. J Cell Physiol 210: 692–702.

36. Nedungadi, T.P., Dutta, M., Bathina, C.S., Caterina, M.J., and Cunningham, J.T. (2012). Expression and distribution of TRPV2 in rat brain. Exp Neurol 237: 223–237.

37. Obukhov, A.G., and Nowycky, M.C. (2005). A cytosolic residue mediates Mg^2+^ block and regulates inward current amplitude of a transient receptor potential channel. J Neurosci 25: 1234–1239.

38. Pearlman, S.M., Serber, Z., and Ferrell, J.E., Jr. (2011). A mechanism for the evolution of phosphorylation sites. Cell 147: 934–946.

39. Peng, G., Lu, W., Li, X., Chen, Y., Zhong, N., Ran, P., and Wang, J. (2010). Expression of store-operated Ca^2+^ entry and transient receptor potential canonical and vanilloid-related proteins in rat distal pulmonary venous smooth muscle. Am J Physiol Lung Cell Mol Physiol 299: L621–630.

40. Samways, D.S., and Egan, T.M. (2011). Calcium-dependent decrease in the single-channel conductance of TRPV1. Pflugers Arch 462: 681–691.

41. Santoni, G., Farfariello, V., Liberati, S., Morelli, M.B., Nabissi, M., Santoni, M., and Amantini, C. (2013). The role of transient receptor potential vanilloid type-2 ion channels in innate and adaptive immune responses. Front Immunol 4: 34.

42. Shibasaki, K., Murayama, N., Ono, K., Ishizaki, Y., and Tominaga, M. (2010). TRPV2 enhances axon outgrowth through its activation by membrane stretch in developing sensory and motor neurons. J Neurosci 30: 4601–4612.

43. Shimosato, G., Amaya, F., Ueda, M., Tanaka, Y., Decosterd, I., and Tanaka, M. (2005). Peripheral inflammation induces up-regulation of TRPV2 expression in rat DRG. Pain 119: 225–232.

44. Siveen, K.S., Nizamuddin, P.B., Uddin, S., Al-Thani, M., Frenneaux, M.P., Janahi, I.A., Steinhoff, M., and Azizi, F. (2020). TRPV2: A Cancer Biomarker and Potential Therapeutic Target. Dis Markers 2020: 8892312.

45. Stokes, A.J., Shimoda, L.M., Koblan-Huberson, M., Adra, C.N., and Turner, H. (2004). A TRPV2-PKA signaling module for transduction of physical stimuli in mast cells. J Exp Med 200: 137–147.

46. Studer, M., and McNaughton, P.A. (2010). Modulation of single-channel properties of TRPV1 by phosphorylation. J Physiol 588: 3743–3756.

47. Sugio, S., Nagasawa, M., Kojima, I., Ishizaki, Y., and Shibasaki, K. (2017). Transient receptor potential vanilloid 2 activation by focal mechanical stimulation requires interaction with the actin cytoskeleton and enhances growth cone motility. FASEB J 31: 1368–1381.

48. Tian, Q., Hu, J., Xie, C., Mei, K., Pham, C., Mo, X., Hepp, R., Soares, S., Nothias, F., Wang, Y., et al. (2019). Recovery from tachyphylaxis of TRPV1 coincides with recycling to the surface membrane. Proc Natl Acad Sci U S A 116: 5170–5175.

49. Voets, T., Nilius, B., Hoefs, S., van der Kemp, A.W., Droogmans, G., Bindels, R.J., and Hoenderop, J.G. (2004). TRPM6 forms the Mg^2+^ influx channel involved in intestinal and renal Mg^2+^ absorption. J Biol Chem 279: 19–25.

50. Wang, Y., Mo, X., Ping, C., Huang, Q., Zhang, H., Xie, C., Zhong, B., Li, D., and Yao, J. (2020). Site-specific contacts enable distinct modes of TRPV1 regulation by the potassium channel Kvbeta1 subunit. J Biol Chem 295: 17337–17348.

51. Yamashiro, K., Sasano, T., Tojo, K., Namekata, I., Kurokawa, J., Sawada, N., Suganami, T., Kamei, Y., Tanaka, H., Tajima, N., et al. (2010). Role of transient receptor potential vanilloid 2 in LPS-induced cytokine production in macrophages. Biochem Biophys Res Commun 398: 284–289.

52. Yang, F., Ma, L., Cao, X., Wang, K., and Zheng, J. (2014). Divalent cations activate TRPV1 through promoting conformational change of the extracellular region. J Gen Physiol 143: 91–103.

53. Yao, J., Liu, B., and Qin, F. (2009). Rapid temperature jump by infrared diode laser irradiation for patch-clamp studies. Biophys J 96: 3611–3619.

54. Yu, L., Xu, L., Xu, M., Wan, B., Yu, L., and Huang, Q. (2011). Role of Mg^2+^ ions in protein kinase phosphorylation: insights from molecular dynamics simulations of ATP-kinase complexes. Molecular Simulation 37: 1143–1150.

55. Zanou, N., Mondin, L., Fuster, C., Seghers, F., Dufour, I., de Clippele, M., Schakman, O., Tajeddine, N., Iwata, Y., Wakabayashi, S., et al. (2015). Osmosensation in TRPV2 dominant negative expressing skeletal muscle fibres. J Physiol 593: 3849–3863.

56. Zhang, Q., Tang, Z., An, R., Ye, L., and Zhong, B. (2020). USP29 maintains the stability of cGAS and promotes cellular antiviral responses and autoimmunity. Cell Res 30: 914–927.

57. Zhang, X., Huang, J., and McNaughton, P.A. (2005). NGF rapidly increases membrane expression of TRPV1 heat-gated ion channels. Embo j 24: 4211–4223.

58. Zubcevic, L., Herzik, M.A., Jr., Chung, B.C., Liu, Z., Lander, G.C., and Lee, S.Y. (2016). Cryo-electron microscopy structure of the TRPV2 ion channel. Nat Struct Mol Biol 23: 180–186.

